# From Hype to Health Check: Critical Evaluation of Drug Response Prediction Models with DrEval

**DOI:** 10.1101/2025.05.26.655288

**Authors:** Judith Bernett, Pascal Iversen, Mario Picciani, Mathias Wilhelm, Katharina Baum, Markus List

## Abstract

**Motivation:** Large-scale drug sensitivity screens have enabled training drug response prediction models based on cancer cell line omics profiles to facilitate personalized medicine. While model performances reported in the literature appear promising, no successful translation to the clinic has been reported.

**Results:** We identify six primary obstacles to this: non-reproducible models, data leakage leading to poor generalization, pseudoreplication, biased evaluation metrics, missing ablation studies, and inconsistent viability data reducing comparability across models. Together, these issues lead to overly optimistic performance estimates of state-of-the-art models and make it challenging to track progress in the field.

To address this, we present DrEval, a pipeline for unbiased and biologically meaningful evaluation of cancer drug response models. It includes baseline and literature models with consistent hyperparameter tuning, statistically sound evaluations, and cross-study benchmarks. DrEval enables ablation studies and publication-ready visualizations. It allows researchers to focus on model development without implementing their own evaluation protocol.

We find that deep learning models barely outperform a naive model predicting the mean drug and cell line effects, while no complex model significantly outperforms properly tuned tree-based ensemble baselines in relevant settings. We advocate making our pipeline a standard benchmark for cancer drug response prediction, ensuring a clinically relevant and robust assessment.

**Availability and implementation:** DrEval consists of a Python package, available on PyPI (drevalpy) and GitHub (github.com/daisybio/drevalpy), and an accompanying nf-core pipeline (github.com/nf-core/drugresponseeval). All data is available on Zenodo (DOI: 10.5281/zenodo.12633909), preprocessing scripts on github.com/daisybio/preprocess_drp_data.

## 1. Introduction

Cancer is a highly heterogeneous class of diseases caused by uncontrolled cell growth and spread. Treatment choice is usually based on broad subtype classification, e.g., by the tissue of origin and a few genetic markers [Liu et al., 2024]. This approach fails to account for the complexity of cancer, leading to unpredictable and inconsistent treatment outcomes [Garraway and Jänne, 2012]. Multi-omics characterizations of tumor cell lines and corresponding large-scale drug response screens have opened up the possibility of linking treatment resistance to the molecular profile of tumor cells. Examples of drug response screens include the Cancer Cell Line Encyclopedia (CCLE) [Barretina et al., 2012], the Cancer Therapeutics Response Portal (CTRP) [Seashore-Ludlow et al., 2015], and Genomics of Drug Sensitivity in Cancer (GDSC) [Iorio et al., 2016], which exposed cultivated tumor cell lines to cancer drugs, measuring cell survival given various concentrations (viability assays). For drug response prediction, metrics summarizing the fitted, typically sigmoidal dose-response relationship curve are used, such as the half-maximal inhibitory/effective concentration (*IC*50/*EC*50) or the area under the curve (*AUC*). Predicting drug response using the unperturbed omics profile of these cultivated cell lines has become an active area of research with strategies ranging from statistical models and network analyses to complex deep learning models [Cortes-Ciriano et al., 2016, Firoozbakht et al., 2022, Partin et al., 2023]. The approaches can be categorized into single-drug models and global approaches. Single-drug models are trained to predict the drug response for a specific drug and thereby only rely on cell line-characterizing features. In contrast, global models are trained across multiple drugs and utilize drug features, such as their chemical properties, theoretically enabling them to predict response values for unseen drugs. While model performance in the literature appears promising, no successful translation of these models to clinical applications that can effectively inform medical decisions has been reported. This gap highlights the need to study and address the challenges that hinder transferable progress in the field [Schätzle et al., 2020].

### 1.1. Challenges in drug response prediction

We identify six primary obstacles to meaningful progress in the drug response modeling field (Figure 1).

**Fig. 1:**
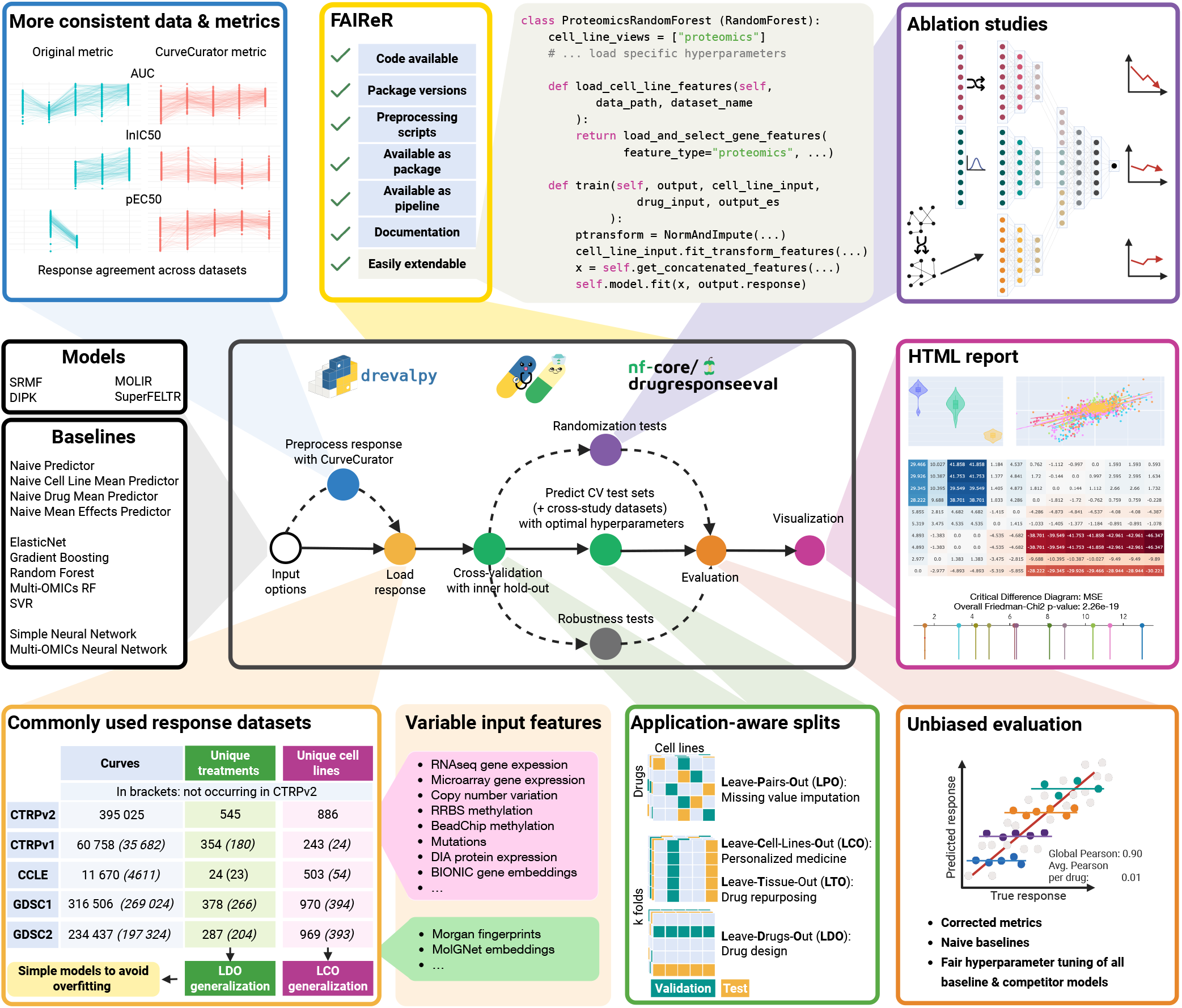
Overview of the DrEval framework. Via input options, implemented state-of-the-art models can be compared against baselines of varying complexity. We address obstacles to progress in the field at each point in our pipeline: Our framework is available on PyPI and nf-core and we follow FAIReR standards for optimal reproducibility. DrEval is easily extendable as demonstrated here with a pseudocode implementation of a proteomics-based random forest. Custom viability data can be preprocessed with CurveCurator, leading to more consistent data and metrics. DrEval supports five widely used datasets with application-aware train/test splits that enable detecting weak generalization. Models are free to use provided or custom cell line– and drug features. The pipeline supports randomization-based ablation studies and performs robust hyperparameter tuning for all models. Evaluation is conducted using meaningful, bias-resistant metrics to avoid inflated results from artifacts such as Simpson’s paradox. All results are compiled into an interactive HTML report. Created in https://BioRender.com.

#### Reproducibility crisis in drug response prediction

Kapoor and Narayanan [2023] postulate that machine learning (ML)-based science is in a reproducibility crisis. They adopt a broader definition of reproducibility, requiring not only computational reproducibility (using the available code and data), but also correct analysis of the data. An analysis by Overbeek et al. [2024] highlights significant barriers to the reusability of drug response models, citing a lack of modular source code, comprehensive preprocessing information, and adequate documentation. Consequently, drug response prediction is highly affected by the reproducibility crisis, and clear standards of reproducible model sharing are needed.

#### Data leakage and weak generalization

Data leakage occurs when information from the test set (inadvertently) influences the training process, leading to optimistically biased risk estimates and comparatively poor generalization performance [Bernett et al., 2024]. One common source of data leakage is incorrect test set design. The data splitting into training, validation, and test set has to fit the intended application of the model, with the test set resembling production data.

For the use case of personalized medicine, where the model needs to generalize to unseen patients, the test set must not contain cell lines used for training or tuning. We refer to this as the leave-cell-line-out (LCO) setting. In contrast, cancer drug repurposing aims to determine whether a drug effective in one cancer type may also be effective in another. This application scenario requires models to generalize across tissue types. Therefore, the test set must contain tissues of origin not seen during training (leave-tissue-out/LTO). In principle, the global models could be applied to drug design. Here, the model must generalize over the chemical input space, and the test set should contain only unseen drugs (leave-drug-out/LDO). Leave random drug-cell line pairs out (LPO) is only warranted when the goal is to evaluate the ability to impute missing values, e.g., for models that guide treatment selection based on a few measured drug responses.

Previous experiments have shown that models especially struggle in the LDO setting and that seemingly good performance measures for LPO and LCO can be reached by predicting the average response of each drug across the training cell lines [Li et al., 2023, Partin et al., 2023]. Developers should, hence, explicitly state for which application scenarios their model is developed and then evaluate its performance in the proper setting against the appropriate baselines.

#### Pseudoreplication

Drug response datasets provide an extensive number of measurements (Figure 1, Table S2). This abundance of data prompts developers to implement complex deep learning models with millions of learnable parameters [Partin et al., 2023]. However, the data points are pseudoreplicated from, e.g., 969 unique cell lines and 287 drugs for GDSC2.

In general, pseudoreplication occurs when multiple measurements from the same unit are treated as if they were from separate independent units [Hurlbert, 1984]. This significantly exacerbates the problem of choosing the right number of features. If, e.g., a model should generalize to an unseen cell line and is trained on the GDSC2 viability data, there are only 969 cell line examples from which it can learn this task. If the number of features and learnable parameters is much larger, the model is likely to overfit by capturing noise rather than genuine drug resistance patterns. Furthermore, statistical testing of model performances can be misdesigned due to pseudoreplication, potentially exaggerating the significance of results.

#### Biased evaluation

Drugs differ considerably in their pharmacologic mode of action, binding affinity, and cellular uptake rate, leading to activity at vastly different concentrations or doses. As a result, the primary source of variance in drug response data, measured by half-maximal concentrations such as *IC*50, arises from differences in the mean drug response across these diverse mechanisms of action of the anti-cancer drugs. This gives rise to a version of Simpson’s paradox during evaluation: a model that simply memorizes each drug’s mean *IC*50 value can explain most of the variance in the data [Li et al., 2023]. However, this success is misleading, as the model fails to capture meaningful drug sensitivity patterns driven by the phenotypic differences in cell lines, thus not learning any generalizable biomedical insights.

Many works also employ a specific set of hyperparameters but do not publish a tuning workflow. In case the hyperparameters are selected based on data dredging (i.e., repeatedly testing different configurations on the test set on which the final metrics are reported until finding ones that yield favorable results), the reported model performance becomes artificially inflated [Scheffer and Herbrich, 1997]. Furthermore, if the baseline methods are not thoroughly tuned, their performance is likely underestimated, making the new model seem superior.

#### Missing ablation studies

To build on the current state-of-the-art, researchers need to be able to assess which modeling components are currently successful [Meyes et al., 2019]. While drug response models grow in complexity and increasingly incorporate multi-modal data, ablation studies (systematic perturbation of model parts) are rarely performed [Partin et al., 2023]. Without proper ablation studies, it is difficult to pinpoint the sources of improvement, making the evaluation of the model’s contributions incomplete.

#### Inconsistency of data and lack of consistent benchmarks

Drug response data is inconsistently preprocessed: Studies apply different normalization techniques, such as min-max scaling or z-score normalization, to the logarithmic *IC*50 target, or binarize responses as sensitive or resistant. Given that the *IC*50 is not distributed bimodally, thresholds for this distinction are often set subjectively. These differences render response metrics (Figure S1) and reported error scores incomparable and make a standardized benchmark with uniformly processed response data a prerequisite for progress. Scientific fields have seen dramatic leaps in progress fueled by standardized benchmarks. Benchmarks like ImageNet [Deng et al., 2009], GLUE [Wang et al., 2018], and CASP [Kryshtafovych et al., 2023] provided rigorous, community-accepted evaluation standards that enabled transparent comparisons and drove algorithmic innovation. In contrast, drug response prediction still lacks a robust, shared benchmark.

### 1.2. The DrEval framework

To address these challenges, we introduce DrEval, a framework for drug response prediction benchmarking (Figure 1). DrEval ensures that evaluations are bias-free, application-oriented, and reproducible. It simplifies the implementation of drug response prediction models, allowing researchers to focus on advancing their modeling innovations by automating standardized evaluation protocols and preprocessing workflows. DrEval includes fair and consistent hyperparameter tuning. Its flexible model interface supports any model type, ranging from statistical models to complex neural networks.

DrEval consists of a standalone Python package available on PyPI as drevalpy and an accompanying Nextflow pipeline, which is part of nf-core [Ewels et al., 2020], a framework for community-curated bioinformatics pipelines. The pipeline enables executing DrEval on large compute clusters in a reproducible environment, guaranteeing reproducibility and scalability. Users automatically benefit from Nextflow features such as execution reports detailing runtime and memory usage. Further, the nf-core initiative promotes collaboration over redundancy by encouraging researchers to jointly develop and maintain best-practice pipelines rather than independently re-implementing similar solutions. In the context of drug response prediction, this consolidates efforts into a single, well-maintained pipeline, reducing fragmentation, fostering reproducibility, and providing a straightforward, standardized workflow that serves as a guideline for implementing, evaluating, and sharing drug response models. DrEval adheres to FAIReR standards: datasets and code used to run the models are findable (F) and accessible (A), the pipeline is executable with conda, Docker, or Singularity, hence, interoperable (I), through the pipeline or the standalone, models and evaluations are runnable and reusable (R) via one command, making all our shown results reproducible (eR) [Tiwari et al., 2023].

### 1.3. Related work

Several benchmark studies assess the relative effectiveness of drug response modeling strategies, focusing on specific components such as gene embeddings [Jia et al., 2023], transcriptome-based vs. marker-based models [Nguyen et al., 2017], dimensionality reduction [Eckhart et al., 2024], pathway-based neural networks [Li et al., 2023], and multi-omics integration [Hauptmann and Kramer, 2023].

Cross-study generalization is examined by Xia et al. [2022] and Partin et al. [2025], who both find generalization to be limited due to overfitting and assay-specific differences (e.g., Syto60 in GDSC1 vs. CellTiter-Glo in other screens).

Efforts to establish fair and consistent benchmarking protocols fall short in key areas. Hauptmann and Kramer [2023] re-implement multi-omics models with standardized preprocessing, hyperparameter tuning, statistical comparisons, and ablation studies. However, their evaluation focuses on classification tasks and excludes the relevant LCO, LTO, and LDO prediction scenarios. In addition, complex models are not compared against simple baseline models, and the implementation does not allow straightforward ways to contribute, limiting future extension and broad adoption in the field. Szalai et al. [2023] highlight inflated performance due to drug-specific mean *IC*50 differences and the fully randomized splitting, and propose bias-corrected evaluation metrics. However, their scope is limited to three basic models trained and tested on the GDSC2 dataset, only using gene expression and mutation features as predictors. Implementation of their bias correction is available as a collection of Jupyter notebooks. They further provide predefined data splits. This, however, constrains the reproducibility and testing of more general settings, which hinders the automation of evaluation and comparison workflows by other researchers. Partin et al. [2025] introduce a Python-based benchmarking package focused on cross-dataset generalization. While being standardized and allowing automation, it requires separate scripts and re-implementations per model. Further, it still lacks support for biologically meaningful data splitting strategies, while only using omics input from DepMap, a cancer data community resource [Tsherniak et al., 2017]. In addition, the package evaluates the provided models in a narrow hyperparameter space with simple parallelization techniques that make automation of larger grid searches on compute clusters difficult. No publication examines the generalization to unseen tissues of origins (LTO setting). Besides, drug response data employed by most studies disregards the variability between replicates.

In contrast, DrEval fits curves using replicate-level data with CurveCurator [Bayer et al., 2023], to allow for consistent, robust estimation of common viability-based drug response measures across datasets (Figure S1). Further, DrEval offers a unified model interface for extensibility and fully supports realistic splitting scenarios that test model applicability in relevant real-world scenarios. It enables unrestricted and custom hyperparameter tuning, evaluates generalization capabilities, and is available as an accompanying Nextflow pipeline for scalability in high-performance computing environments. Instead of delivering a state-of-the-art snapshot, DrEval is designed as a continuously extensible framework supporting systematic experimentation, facilitating integration of new models and datasets, and promoting long-term reproducibility and collaboration.

## 2. Methods

### 2.1. Model interface and integration in DrEval

DrEval defines a model as a function that maps a cell line and drug pair to a continuous response value (regression). The inputs characterize the cell lines (e.g., baseline omics measurements) and drugs (e.g., fingerprints) and are flexible across models. Each model defines its own feature loading. The preprocessing of the input features is seen as part of the modeling process and can be defined uniquely for each model. To ensure a fair comparison between models, DrEval fixes the output across all compared models. Supported outputs are *lnIC*50, *IC*50, *EC*50, *pEC*50 (negative *log*10 of the *EC*50), and *AUC*. Additionally, arbitrary measures can be used when providing custom datasets, but at the moment, we only support regression models. If not specified otherwise, we predicted *lnIC*50 for this study as it is most commonly used by state-of-the-art models [Partin et al., 2023].

DrEval employs a flexible model interface, supporting any models that admit `train` and `predict` procedures. Developers must implement their feature loading and, optionally, a set of hyperparameters for model tuning. Accordingly, DrEval allows for any modeling strategy, such as statistical models, ML, or algorithmic approaches, and any input feature type.

### 2.2. Implemented baseline and literature models

An overview of all implemented models can be found in Table S1. We re-implemented several models from the literature, covering a range of different strategies: SRMF (matrix factorization) [Wang et al., 2017] and DIPK (complex deep neural network with gene and drug features) [Li et al., 2024] are included as global models. We also include single-drug models that are fitted and tuned per drug, respectively, by adapting the classifiers MOLI (most cited simple neural network, multi-omics late integration) [Sharifi-Noghabi et al., 2019] and SuperFELT (refinement of MOLI) [Park et al., 2021] to regression. Researchers can employ these models as baselines to compare against their work, eliminating the need to tediously re-implement the models from disparate source codes.

Furthermore, we offer global, gene expression– and drug fingerprint-based baselines, such as a fully connected neural network (early integration) and various classical ML models (Elastic Net, Random Forest, Gradient Boosting, Support Vector Regressor). The neural network and random forest are also available as multi-omics version (additionally employing copy number variation, methylation, and mutation features). The hyperparameter search grids for all models are listed in Table S3. DrEval includes a set of naive predictors based solely on response statistics to assess whether complex models truly capture interactions or simply exploit marginal effects. These consist of the *NaivePredictor*, which returns the average response of the training set, the *NaiveDrugMeanPredictor*, which returns the average response for each drug, and the *NaiveCellLineMeanPredictor*, which predicts the average response per cell line. The *NaiveMeanEffectsPredictor* combines both sources of variation and predicts responses as the sum of the overall mean with cell line and drug effects

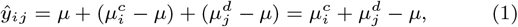

where *μ* is the NaivePredictor, 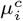 is the mean for cell line *i*, and 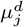 is the mean for drug *j*. If predictions are made for a drug or cell line that was not observed, we set 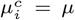 or 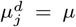, respectively. Finally, the *NaiveTissueMeanPredictor* predicts the mean response of all training cell lines that belong to the same tissue of origin as the cell line, for which predictions are made. Despite their simplicity, these models can appear to perform well when variance in the data is attributable to systematic differences in mean drug response.

### 2.3. Data splitting and fair hyperparameter tuning

DrEval employs a k-fold cross-validation schema incorporating an inner holdout validation process for model tuning. An early-stopping-validation set is split from the remaining train set for models employing an early stopping mechanism (DIPK, MOLIR, SuperFELTR, Simple / MultiOmicsNeuralNetwork). For the LCO, LDO, and LTO settings, we ensure that all splits are disjoint for cell lines, drugs, and tissues, respectively. In the LPO setting, we only ensure that replicate experiments involving the same drug-cell line pair are contained within the same data subset. We enforce a fair setting by tuning all baselines of a benchmark in addition to the tested model so that performance gains are due to genuine model enhancements rather than suboptimal baseline configurations.

### 2.4. Benchmark data

#### Cell viability data

Raw dose-response data was downloaded from five publicly available, commonly used datasets: CCLE (Supplement of Barretina et al. [2012]), CTRPv1/v2 (CTD^2^ Data Portal), and GDSC1/2 (cancerrxgene website). Dataset sizes are shown in Figure 1. All preprocessing scripts are available in the accompanying GitHub repository. All cell line names were mapped to Cellosaurus CVCL accessions and all drug names to PubChem CIDs (when available).

To ensure consistency and reduce preprocessing discrepancies, all datasets are reprocessed in a standardized workflow made available via `drevalpy` (Figure S1). Hence, we also support fitting custom data, facilitating cross-study predictions through compatibility with the provided datasets and future dataset contributions.

Instead of aggregating replicates prior to normalization and curve fitting, as is the standard practice, DrEval includes replicate variability into quality control measures. This source of experimental variability is often overlooked when aggregating replicates prior to fitting, which leads to inaccurate or misleading drug response measures in case of large discrepancies between replicates. Therefore, raw measurements are normalized per replicate by dividing by the control (no-drug) measurement, yielding viability values from 0 to 1; these are then used by CurveCurator [Bayer et al., 2023] to fit a single model across replicates for each drug–cell line pair. This allows CurveCurator to include residual differences at each measurement point for more robust estimation of quality measures, compared to simply taking the mean or median of the replicates at each measurement point. For the CCLE dataset, only aggregated data was available, which had already been normalized, so it was treated as a single replicate.

CurveCurator calculates p-values using a recalibrated F-statistic for each curve fit, measuring how well the data conforms to a sigmoidal curve, which is the biologically expected phenotypic response to increasing drug dosages. Since different datasets test varying dosage ranges for specific drugs, curve fitting and p-value calculations were conducted within each dosage group to ensure fair comparisons. Finally, *EC*50 values falling outside the measured dosage range per drug and *IC*50 values deviating by more than one order of magnitude from the measured range were considered invalid. The datasets can be filtered for quality using the statistical measures provided by CurveCurator, minimizing inter-dataset batch effects. This leaves experimental protocols and the choice of viability assays as the primary potential sources of systematic shifts. If not stated differently, we did not apply any quality filter in our benchmark experiments to maintain comparability to previous studies and avoid data loss.

#### Cell line features

Omics screens corresponding to the five employed drug response screens are available from various sources, overlapping at varying proportions (Table S2). We supply two sources of gene expression and methylation data because they better complement CCLE/CTRPv1/CTRPv2 or GDSC1/GDSC2. For the CCLE and CTRP screens, multiple gene expression datasets (microarray, RNA-seq) exist with varying measures and preprocessing, leading to inconsistencies. To ensure reproducibility, we reprocessed the raw RNA-seq data from Ghandi et al. [2019] (PRJNA523380) using the nf-core RNA-seq pipeline [Patel et al., 2023] (STAR for alignment [Dobin et al., 2013], Salmon for quantification [Patro et al., 2017], version 3.10.1). We supply the resulting TPM values. As a second source complementing the GDSC screens, we obtained RMA-normalized microarray expression data from the GDSC Data Portal. Methylation BeadChip data for GDSC (preprocessed beta values for all CpG islands) was also obtained from the GDSC Data Portal. Further, methylation RRBS data for promoter CpG clusters were downloaded from DepMap (Release: Methylation (RRBS)) [Tsherniak et al., 2017], which has a larger overlap with the CCLE and CTRP screens.

Mutation and copy number variation (CNV) data was downloaded from Cell Model Passports [van der Meer et al., 2019] and can be combined with all response screens. The mutation data is binary and was filtered to only contain coding mutations that are not silent. CNV is represented as GISTIC scores derived from Affymetrix SNP6.0 array data (integers ranging from *−*2: high-level deletion to 2: high-level amplification). Proteomics data was obtained from a DIA screen by Gonçalves et al. [2022]. The raw data (PRIDE: PXD030304) was filtered for global Q values *≤* 0.01 and proteotypic peptides. Protein quantities were calculated using the MaxLFQ algorithm of the DIA-NN package (version 1.0.1) [Demichev et al., 2020]. We offer to filter genes with gene sets such as anti-cancer drug target genes curated by GDSC, the landmark genes used by the L1000 assay [Subramanian et al., 2017], or user-provided sets.

#### Drug features

We use Morgan fingerprints generated from SMILES with RDKit [2025] for drug representation. GDSC1 and GDSC2 already provide SMILES. For the other datasets, they were downloaded from PubChem [Kim et al., 2025]. DIPK uses 768-dimensional drug encodings generated using the pre-trained graph neural network-based molecule encoder MolGNet [Li et al., 2021] and 512-dimensional interactome features computed by averaging the embeddings of the top 256 highly expressed gene embeddings extracted from BIONIC [Forster et al., 2022], a graph autoencoder trained on multiple gene interaction networks. We adapted DIPK’s preprocessing scripts to generate missing MolGNet features from SMILES.

#### Tissue information

We obtained disease annotation for our LTO setting from Cellosaurus (Release 52) and enriched it with DepMap (22Q2) sample information if Cellosaurus disease information was missing. We then applied a curated tissue synonym dictionary to map specific diseases to broader, biologically meaningful tissue-of-origin categories. Finally, we manually overwrote mappings for misclassified or ambiguous cell lines based on verified external sources (e.g., ATCC, NCI).

### 2.5. Ablation studies, robustness tests, and cross-study prediction

DrEval supports feature ablation via modular input permutations. Models must declare their input feature modalities (e.g., gene expression, mutation), which allows independent permutation of each view. For instance, if a model uses gene expression and mutation data, DrEval fits variants where one modality is permuted across cell lines (e.g., shuffling gene expression profiles), removing any possible connection between this modality and the response. A performance drop indicates the importance of that modality. To determine whether a model relies on informative biological signals rather than simple summary statistics (such as mean and variance), DrEval also implements summary statistics-invariant randomizations. We generate synthetic features drawn from a normal distribution preserving the original mean and standard deviations for tabular features. For graph inputs, edges are shuffled while preserving node degrees. This shows whether a model learns from specific biological signals or fits simple statistical attributes, for example, the overall mean gene expression of a cell line.

For robustness tests, DrEval refits models with different initializations to evaluate the performance stability. We further implement cross-study predictions to assess cross-dataset generalization performance. In our LPO setting, no cross-study predictions are made for drug-cell line combinations present in the training dataset. In the LCO, LTO and LDO settings, cross-study predictions for cell lines, tissues or drugs found in the training dataset are excluded.

### 2.6. Evaluation and visualization

We compute the following metrics between the true and predicted responses: mean squared error (MSE), root MSE (RMSE), mean absolute error (MAE), the coefficient of determination (*R*^2^), Pearson, Spearman, and Kendall correlation. Metrics are calculated separately for each cross-validation fold based on all test set predictions. In addition, we assess the *R*^2^ and correlation metrics stratified by drug and by cell line, aggregating predictions across all folds. We also define normalized *R*^2^ and correlation coefficients by subtracting the predictions of the NaiveMeanEffectsPredictor from both predicted and true responses before evaluation. This removes any variance due to average drug and cell line effects. The remaining variance includes both noise and differential drug response, which is the variation driven by interactions between molecular cell line features and drug properties. These effects are clinically relevant, as they reflect mechanisms of sensitivity and resistance of cancer cells. We further compute effect sizes by calculating strictly standardized mean differences for the *R*^2^ and MSE. We assess statistical significance using the Friedman test on MSE-based model ranks across CV folds. If significant, a Conover post-hoc test identifies which model pairs differ. For both tests, we set a significance level of 0.05. We present the results in an automatically generated HTML report with interactive plots. An example is available on GitHub.^1^ All underlying evaluation metrics, ground truth values, and predictions are also provided as CSV files for further analysis.

### 2.7. Benchmarking setup

We benchmark all models using 10-fold cross-validation in the implemented LPO, LCO, LTO, and LDO settings. To ensure fairness, all baseline and test models are tuned based on the validation RMSE using the same cross-validation folds, and performance is assessed using MSE, *R*^2^, Pearson correlation, and their normalized variants. The hyperparameter spaces are inferred from the relevant publications or set to balance run time and search space size (Table S3). Since it is the largest available dataset, the primary training and evaluation dataset is CTRPv2 with CurveCurator-processed responses. For all gene expression-based models, we use the RNAseq gene expression data. For models requiring methylation data, we use the RRBS methylation data. Our cross-study evaluations assess generalization to the CurveCurator-processed responses of CTRPv1 and CCLE using the same input screens.

## 3. Results

To create a living benchmark for drug response prediction, we present DrEval, a systematic evaluation framework focused on reproducibility and extensibility (Figure 1). While allowing flexible model design and feature engineering, we enforce strict management of the predictor, preventing target leakage and ensuring bias-free evaluation. We conducted an extensive benchmark across multiple model strategies and application settings using this framework.

### 3.1. Models learn limited biological signal beyond drug means

An overview of our benchmark results is shown in Figure 2) (the full results with all metrics and standard errors are listed in Tables S4-5). About half of the tested models do not significantly outperform the NaiveMeanEffectsPredictor (Figure S2). For those that do, the *R*^2^-gain over the naive model is smaller in the unnormalized than in the normalized case due to the reduced total variance in the normalized setting, which makes the explained variance proportionally greater. In LCO, the setting relevant to personalized medicine applications (generalizing to unseen cell lines), no model surpasses a tuned Random Forest.

**Fig. 2:**
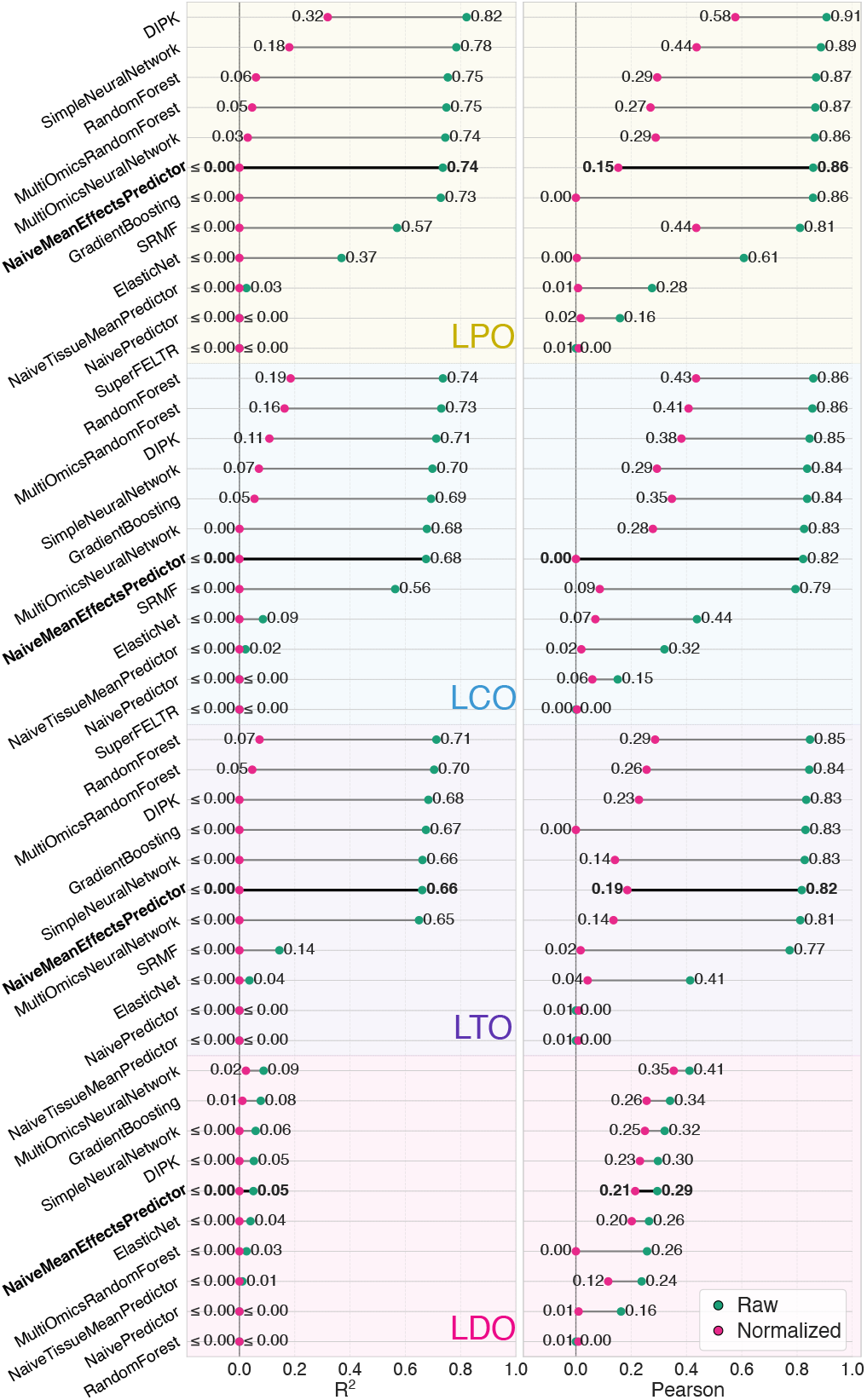
Comparison of model performance for the different test set designs (LPO, LCO, LTO, LDO). We report the mean *R*^2^ and Pearson’s correlation (green) between predicted and ground truth *lnIC*50 over the cross validation folds and normalized *R*^2^ and normalized Pearson’s correlation (pink). The normalized metrics are derived by subtracting the predictions of the NaiveMeanEffectsPredictor from the true and predicted values and then recalculating the *R*^2^ or Pearson’s correlation. The bold NaiveMeanEffectsPredictor represents the best performance without utilizing any features, by exploiting cell line and drug biases. SRMF and SuperFELTR are omitted from LDO as they can not predict unseen drugs. Negative (normalized) *R*^2^ values are clipped at zero.

Normalized metrics and drug– and cell-line-specific evaluations reveal that the drug mean variation dominates the explained variance of drug response models (Tables S4-5). For instance, while DIPK achieves an overall Pearson correlation of 0.91 in LPO, the per-drug correlation drops to 0.56, whereas the per-cell line correlation remains nearly unchanged (0.89). The discrepancy between overall and per-drug performance arises from Simpson’s paradox, as illustrated in Figures 3, and S3 for LDO. By removing the mean effects of drugs and cell lines prior to evaluation, we show that current drug response models capture only a limited fraction of the biologically relevant variation. Our series of naive predictors shows how models can appear effective by memorizing dataset-specific effects and allows us to assess the relative impact of these factors on predictive performance (Table 1). Most of the explainable variation in drug response is driven by the drug identity. Smaller contributions stem from cell line biases and tissues identities which can be learned implicitly by models. In the LCO experiments, DIPK and the Random Forest can only explain 11% and 19% of the differential drug sensitivity (normalized *R*^2^), respectively. These results indicate that current models lack the accuracy required for clinical or translational use.

**Table 1.**
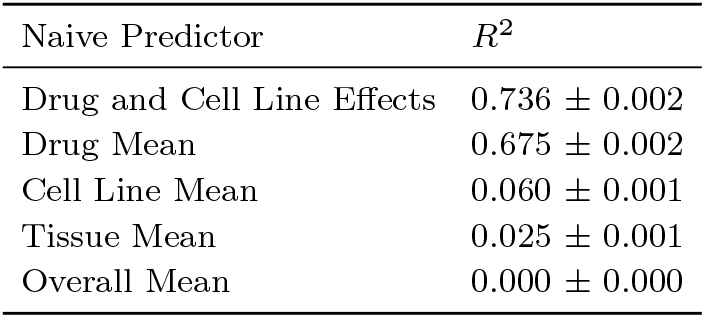
Mean *R*^2^ and standard error across cross-validation folds for naive baseline models in the LPO setting. Overall Mean: *NaivePredictor*, Drug and Cell Line Effects: *NaiveMeanEffectsPredictor*.

| Naive Predictor | $R^2$ |
| --- | --- |
| Drug and Cell Line Effects | $0.736 \pm 0.002$ |
| Drug Mean | $0.675 \pm 0.002$ |
| Cell Line Mean | $0.060 \pm 0.001$ |
| Tissue Mean | $0.025 \pm 0.001$ |
| Overall Mean | $0.000 \pm 0.000$ |

**Fig. 3:**
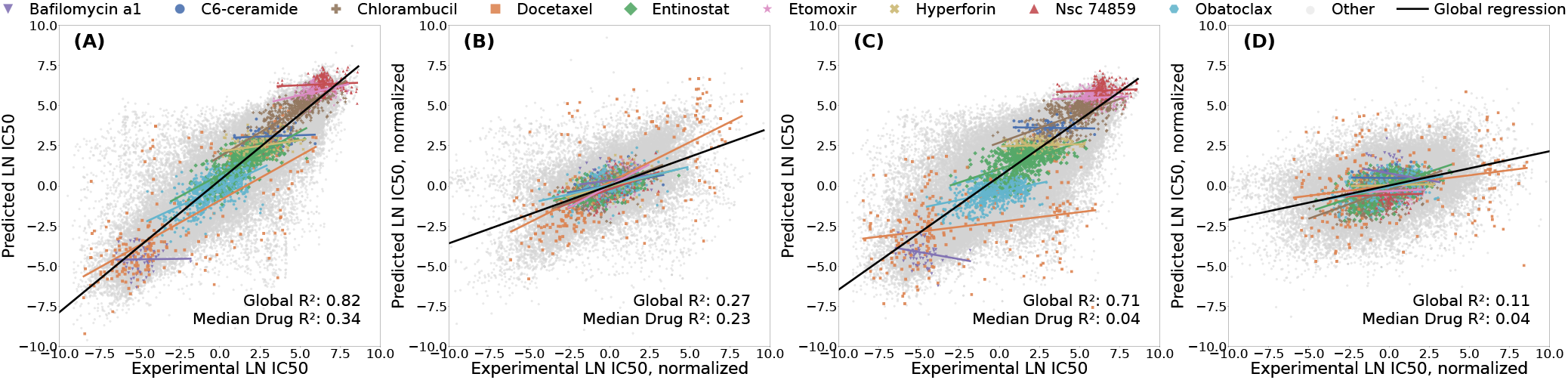
Simpson’s paradox in drug response prediction. (A) Predicted *lnIC*50 vs. ground truth for the DIPK model under leave-pair-out cross-validation. The apparent correlation is largely driven by differences in mean drug potency. (B) After subtracting drug and cell line mean effects, only weak signal remains, indicating limited learning of differential response beyond remembering mean cell line and drug responses. (C) Non-normalized coefficients of determinations are lower under leave-cell-line-out cross-validation. (D) After subtracting drug means for the leave-cell-line-out results, minimal predictive signal remains.

Although DIPK performs comparatively well in the LPO setting, it is the most computationally expensive model. Other models achieve comparable performance with far less training time (Table S6). This highlights the importance of benchmarking against simpler models to ensure that increased complexity translates into meaningful performance gains, given the increased resource consumption.

### 3.2. Models do not generalize to unseen drugs

For the LDO setting, where the task is to predict responses for unseen drugs, no model significantly outperforms the NaiveMeanEffectsPredictor (Figure 4). The near-zero *R*^2^ values indicate that only a negligible amount of the response variance is explained (Figure 2). The tested models fail to learn the relation between molecular drug features and pharmacological effects and do not generalize over the chemical space, likely due to the limited number of drugs in the screens and the high complexity of the structure-activity relationship.

**Fig. 4:**
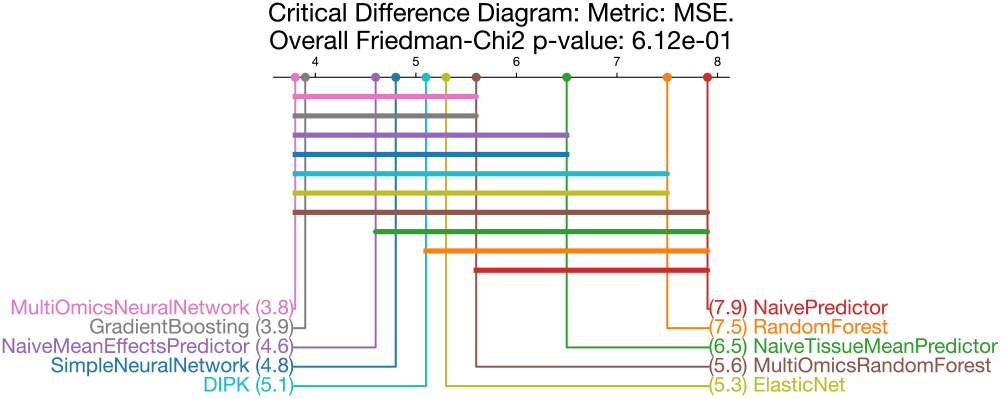
Critical difference diagram for the LDO setting using an MSE-based ranking in the cross-validation folds. For each model, we draw an individual horizontal bar, connecting it to all other models from which it does not differ significantly. While the MultiOmicsNeuralNetwork significantly outperforms the NaiveTissueMeanPredictor, RandomForest and NaivePredictor, it falls short of significantly surpassing the NaiveMeanEffectsPredictor baseline. LPO, LCO, and LTO diagrams in Figure S2.

### 3.3. Weak generalization beyond tissue context

We evaluate the models’ ability to generalize to an unseen tissue of origin in the leave-tissue-out (LTO) setting. LTO better reflects a model’s ability to transfer mechanistic signals across biological contexts, instead of relying on tissue-specific mean differences seen during training. In the unnormalized performance metrics, only a small decrease is observed compared to the LCO setting, as these metrics are primarily dominated by the drug-specific effects. However, when using normalized metrics, we observe a substantial drop in performance (e.g., Random Forest Pearson correlation from 0.43 to 0.29). This indicates that the LCO models have implicitly modeled tissue-identities and that robust generalization to unseen tissues (required for drug repurposing) remains challenging, even when the model hyperparameters are tuned towards it.

### 3.4. Weak generalization across datasets

Cross-study prediction performances are considerably lower compared to within-study performance (Figure 5, Table S7). These observations are consistent over different response metrics (Table S8: comparison *lnIC*50, *AUC, pEC*50). The results indicate that, despite harmonizing the response data using CurveCurator, substantial technical differences between the assays persist, causing weak generalization across datasets.

**Fig. 5:**
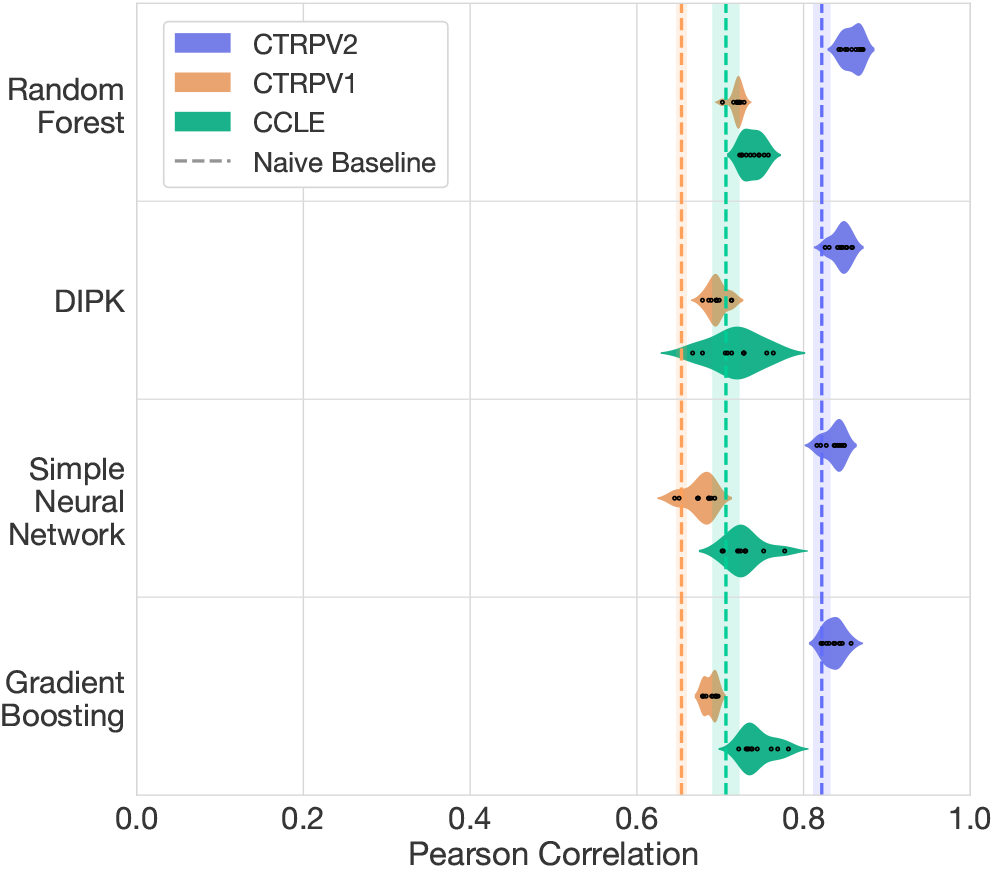
Pearson Correlation of model predictions vs. ground truth *lnIC*50 values for various models. All models are trained on CTRPv2 and tested on unseen cell lines (LCO) of CTRPv2, CTRPv1 and CCLE. Each correlation value corresponds to a single fold of cross-validation. Dashed lines are the mean metrics of the NaiveMeanEffectPredictor colored by the respective test dataset. Shading represents the standard deviation across cross-validation folds.

Due to the limited overlap between RNAseq and RRBS methylation data with GDSC1 and GDSC2, these datasets are typically paired with microarray-based gene expression and BeadChip methylation data (Table S2). This represents a substantial distribution shift in the input features, leading to a further drop in performance (Table S7) when models are applied without accounting for these differences.

### 3.5. Validating model component utility requires ablation studies

We conduct an ablation study to assess the contribution of each data modality by randomly permuting or invariantly randomizing all features of an individual data modality (Figure 6, Table S9). We chose DIPK since it was the best performing multi-modal model and a Multi-Omics Random Forest as a baseline. DIPK relies on gene expression and BIONIC features for cell line representation, and MolGNet encodings for drug representation. The Multi-omics Random Forest is an extension of the gene expression-based Random Forest that additionally incorporates copy number variation, methylation, and mutation data.

**Fig. 6:**
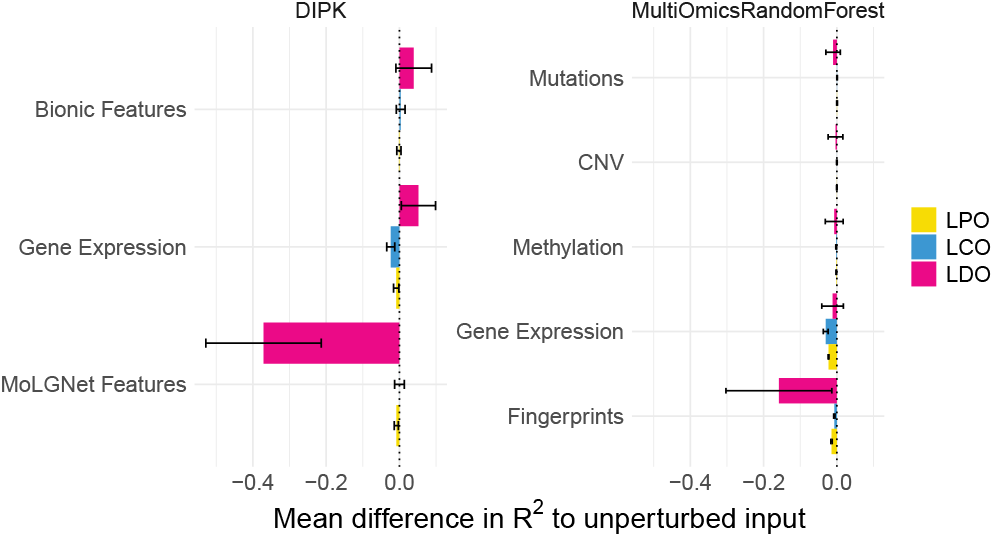
Mean ablation study results of the Multi-omics Random Forest and DIPK. One omics input is randomized at a time, such that a drug/cell line receives a random feature of another drug/cell line. Models are retrained from scratch with the randomized features. The *R*^2^ of the unperturbed models are subtracted from the *R*^2^ of the ablated models for each cross-validation fold: negative values indicate that the perturbation decreased model performance. Error bars are 95% confidence intervals over the cross-validation folds.

For the tested models, no noteworthy differences exist between permutation and invariant randomization (Table S9). Only gene expression and fingerprint/MolGNet randomization results in considerable performance drops. In the LPO setting, both inputs contribute equally to performance, while in the LCO setting, permutations of the gene expression input have a greater impact than permuting the drug input. The drug feature permutation causes the strongest performance drop in the LDO setting, where the model relies on molecular features to extrapolate to unseen drugs. In the LPO and LCO settings, model performance is more dependent on capturing cell line characteristics, which are better represented by gene expression profiles. The Multi-omics Random Forest may have largely ignored the additional omics, as randomizing them did not result in a considerable performance drop. For DIPK, the drug means, and the cell-line specific effects extracted from the gene expression sufficed to reach good performance scores in the LPO and LCO setting, while the BIONIC module did not contribute to the predictive power.

These findings highlight the importance of ablation studies in determining whether integrating additional features or model components provides any benefit. Without careful assessment, added features may introduce noise and exacerbate the curse of dimensionality, especially due to the previously mentioned issue of pseudoreplication.

### 3.6. Community contribution and extensibility: integrating proteomics data

To encourage community-driven benchmarking and reduce duplication of effort, DrEval is designed to be easily extendable. We showcase this by implementing a Random Forest that takes proteomics instead of gene expression data as cell line feature input (pseudocode in Figure 1). Because most methods can be inherited from the Random Forest implementation, only a few lines of code need to be adapted. Once added, the model can immediately be run using the standalone Python package. If contributed via pull request and included in a new release, it becomes available in the Nextflow pipeline, supporting scalable evaluation on large compute clusters. The performance of the proteomics-based model is comparable to the gene expression-based version (Table S10). Researchers can also integrate entirely new model architectures into DrEval with minimal code, facilitating rapid testing and model design under standardized, reproducible conditions.

## 4. Conclusion

Although many cancer drug response prediction models have been proposed, none have been translated into use in clinical practice or drug development. This lack of progress stems from several persistent problems, which DrEval explicitly targets.

First, many models are released without attention to reproducibility, testing, and documentation. A model that cannot be used or re-trained easily will not be applied or refined by the community. DrEval provides a fully versioned Python package and an nf-core pipeline. These come with automated testing, modular design, and continuous integration, ensuring both computational and general reproducibility. We implement state-of-the-art models for comparisons and make contributing easy. Second, in many cases, improper test set design leads to data leakage, and performance metrics are often inflated by drug and cell line mean effects. DrEval enforces strict, use-case-motivated splitting strategies, with normalized metrics to disentangle meaningful prediction from trivial mean effect memorization and overfitting. Third, datasets may appear large due to repeated measurements of the same cell lines and drugs, but this pseudoreplication inflates the apparent sample size and can mislead model design toward complex deep learning models. DrEval includes straightforward statistical and ML baselines, making it clear when added complexity yields meaningful gains. Fourth, many papers report inflated results by tuning only their proposed model. DrEval uses nested cross-validation with tuning for all models and simple baselines, ensuring fair and unbiased comparisons. Fifth, ablation studies are essential to identify which components contribute to performance as models become more multimodal. DrEval provides built-in standardized support for this. Finally, inconsistent preprocessing of drug response data across studies leads to distributional shifts that hinder cross-study training. DrEval addresses this by providing standardized and improved preprocessing workflows and harmonized datasets. This enables research towards generalizability.

Using our pipeline, we find that, currently, complex models barely outperform a naive algorithm predicting the mean drug and cell line effects. In line with this, we demonstrate how commonly used metrics are distorted through Simpson’s paradox. Simple tree-based models perform best in the LCO setting, which is relevant for personalized medicine. This aligns with previous findings [Partin et al., 2023, Szalai et al., 2023]. All models explain less than 20% of the drug sensitivity variance when predicting unseen cell lines (*R*^2^ normalized *<* 0.20). This performance drops further when predicting cell lines of unseen tissues of origin (LTO, *R*^2^ normalized *<* 0.08), a setting relevant for cancer drug repurposing. All tested models fail to predict the effect of an unseen drug (LDO, *R*^2^ *≈* 0). Our ablation studies show that integrating more modalities and more advanced gene or drug representations has little effect, with gene expression being the only source of predictive signals in LPO and LCO. Drug representations matter only in the LDO setting, where the models barely explain any variance. Despite our uniform response data processing, models do not generalize well across datasets. This shows that the technical differences between the screens are substantial, complicating the formation of a large, joined dataset more suitable for deep learning. Possible solutions include establishing standard assays for future screens or solving the problem of dataset integration computationally. In summary, drug response prediction remains a largely unresolved challenge. Despite high reported correlations in the literature, which are inflated due to flawed evaluation, models fail to generalize in realistic settings. Our work highlights these gaps, and our pipeline offers a concrete path forward through rigorous benchmarking, reproducible evaluation, and fast integration of new ideas.

Our study inherits the general limitations of cell line viability-based drug response prediction in cancer cell lines: Predicting drug response from baseline omics is inherently more challenging than doing so with post-treatment molecular profiles, which directly reflects a drug’s effects and enables more informed inferences about its mode of action and similarity to other compounds. Another major unresolved issue is the gap between in vitro measurements and in vivo outcomes [Partin et al., 2023]. Further, even if viability could be predicted accurately, this would not directly translate into valid treatment recommendations. *IC*50 and *EC*50 values are not directly comparable across drugs because different drugs operate at different concentrations or doses. Additionally, viability does not account for the relative toxicity to healthy tissue (where a measure related to side effects or general toxicity, like the LD50, is needed, which response screens cannot determine) [Prasse et al., 2022].

Another limitation is dataset size. With only hundreds of drugs and cell lines, current datasets are too small to capture the complex biochemical effects of molecules on diverse cellular systems. This limitation can most likely not be addressed through better modeling alone and calls for larger, standardized screening efforts.

In the future, we aim to continuously integrate new models to ensure DrEval remains aligned with state-of-the-art developments and encourage researchers to develop their models directly in DrEval. This turns the benchmark into a continually evolving resource that tracks meaningful progress in the field. Since DrEval supports all possible input types, it can be used to study data from proteomics or epigenomics studies, biological network data, and drug representations. In line with this, we plan to support classification strategies, with particular attention to how sensitivity labels are defined, as current practices often rely on thresholding strategies that may not reflect biological or clinical relevance. We further want to add xenograft or patient-derived benchmark data to assess the generalization ability to more clinically relevant settings. Additionally, we plan to include metadata in our evaluations to also account for tissue-, sex-, or age-specific effects.

## 6. Author contributions statement

J.B. (Conceptualization [equal], Data curation [equal], Formal analysis [equal], Investigation [equal], Methodology [equal], Software [equal], Validation [equal], Visualization [equal], Writing – Original Draft [equal], Writing – Review & Editing [equal]), P.I. (Conceptualization [equal], Data curation [equal], Formal analysis [equal], Investigation [equal], Methodology [equal], Software [equal], Validation [equal], Visualization [equal], Writing – Original Draft [equal], Writing – Review & Editing [equal]), M.P. (Conceptualization [supporting], Data curation [supporting], Formal analysis [supporting], Software [supporting], Writing – Original Draft [supporting], Writing – Review & Editing [equal]), M.W. (Conceptualization [supporting], Funding acquisition [equal], Resources [equal], Supervision [supporting], Writing – Review & Editing [equal]), K.B. (Conceptualization [equal], Funding acquisition [equal], Project administration [equal], Resources [equal], Supervision [equal], Writing – Review & Editing [equal]), M.L. (Conceptualization [equal], Funding acquisition [equal], Project administration [equal], Resources [equal], Supervision [equal], Writing – Review & Editing [equal]).

## 7. Acknowledgments

J.B., M.P., M.W., and M.L. were supported by the German Federal Ministry of Education and Research (BMBF) within the framework of the CompLS funding concept [031L0305A (DROP2AI)]. Funded by the Deutsche Forschungsgemeinschaft (DFG, German Research Foundation) [422216132]. The results published here are partially based upon data generated by the Cancer Target Discovery and Development (CTD^2^) Network^2^ established by the National Cancer Institute’s Center for Cancer Genomics. We thank Jonah Reiner for his help with the integration of the DIPK model.

## Footnotes

1 https://dilis-lab.github.io/drevalpy-report/

2 https://www.cancer.gov/ccg/research/functional-genomics/ctd2

