## Supplement for "From Hype to Health Check: Critical Evaluation of Drug Response Prediction Models with DrEval"

### DrEval

Supplementary information

#### 1 Supplementary Material and Methods

Table S1: Overview of the models implemented in DrEval

| Model | Baseline / Published | Global / Single Drug | Description |
|---|---|---|---|
| NaivePredictor | Baseline Method | Global Model | Predicts the mean response of all drugs in the training set. |
| NaiveCellLine-MeanPredictor | Baseline Method | Global Model | Predicts the mean response of a cell line in the training set. |
| NaiveTissue-MeanPredictor | Baseline Method | Global Model | Predicts the mean response of a tissue of origin in the training set. |
| NaiveDrug-MeanPredictor | Baseline Method | Global Model | Predicts the mean response of a drug in the training set. |
| NaiveMean-EffectPredictor | Baseline Method | Global Model | Predicts mean of dataset + mean of cell line + mean of drug |
| ElasticNet | Baseline Method | Global Model | Fits an Sklearn Elastic Net on gene expression data and drug fingerprints. |
| Single-Drug Elastic Net | Baseline Method | Single-Drug Model | Fits an Sklearn Elastic Net on gene expression data for each drug separately. |
| Single-Drug Proteomics Elastic Net | Baseline Method | Single-Drug Model | Fits an Sklearn Elastic Net on proteomics data for each drug separately. |
| Gradient-Boosting | Baseline Method | Global Model | Fits an Sklearn Histogram-based Gradient Boosting Regression Tree on gene expression data and drug fingerprints. |
| RandomForest | Baseline Method | Global Model | Fits an Sklearn Random Forest Regressor on gene expression data and drug fingerprints. |
| MultiOmics-RandomForest | Baseline Method | Global Model | Fits an Sklearn Random Forest Regressor on gene expression, methylation, mutation, copy number variation data, and drug fingerprints (concatenated matrix). The dimensionality of the methylation data is reduced with a PCA to the first 100 components before it is fed to the model. |

| | | | |
|---|---|---|---|
| SingleDrug-RandomForest | Baseline Method | Single-Drug Model | Fits an Sklearn Random Forest Regressor on gene expression data for each drug separately. |
| SVR | Baseline Method | Global Model | Fits an Sklearn Support Vector Regressor on gene expression data and drug fingerprints. |
| Simple-NeuralNetwork | Baseline Method | Global Model | Fits a feedforward neural network on gene expression and drug fingerprints with 3 layers + dropout |
| MultiOmics-NeuralNetwork | Baseline Method | Global Model | Fits a feedforward neural network on gene expression, methylation, mutation, copy number variation data, and drug fingerprints, 3 layers + dropout. The dimensionality of the methylation data is PCA-reduced before it is fed to the model. |
| SRMF | Published Model | Global Model | Similarity Regularization Matrix Factorization model by Wang et al. on gene expression data and drug fingerprints. Similarities to all other drugs/cell lines represent each drug and cell line and are mapped into a shared latent low-dimensional space from which responses are predicted. |
| MOLIR | Published Model | Single-Drug Model | Regression extension of MOLI: multi-omics late integration deep neural network by Sharifi-Noghabi et al. Takes mutation, copy number variation, and gene expression data as input. MOLI reduces the dimensionality of each omics type with a hidden layer, concatenates them into one representation, and optimizes this representation via a combined cost function consisting of a triplet loss and a binary cross-entropy loss. We implemented a regression adaption with MSE loss and an adapted triplet loss for regression. |
| SuperFELTR | Published Model | Single-Drug Model | Regression extension of SuperFELT: supervised feature extraction learning using triplet loss for drug response by Park et al. Very similar to MOLI(R). In MOLI(R), encoders and the classifier were trained jointly. SuperFELT(R) trains them independently. |
| DIPK | Published Model | Global Model | Deep Neural Network Integrating Prior Knowledge from Li et al. Uses gene interaction relationships (encoded by a graph auto-encoder), gene expression profiles (encoded by a denoising auto-encoder), and molecular topologies (encoded by MolGNet). Those features are integrated using multi-head attention layers. |

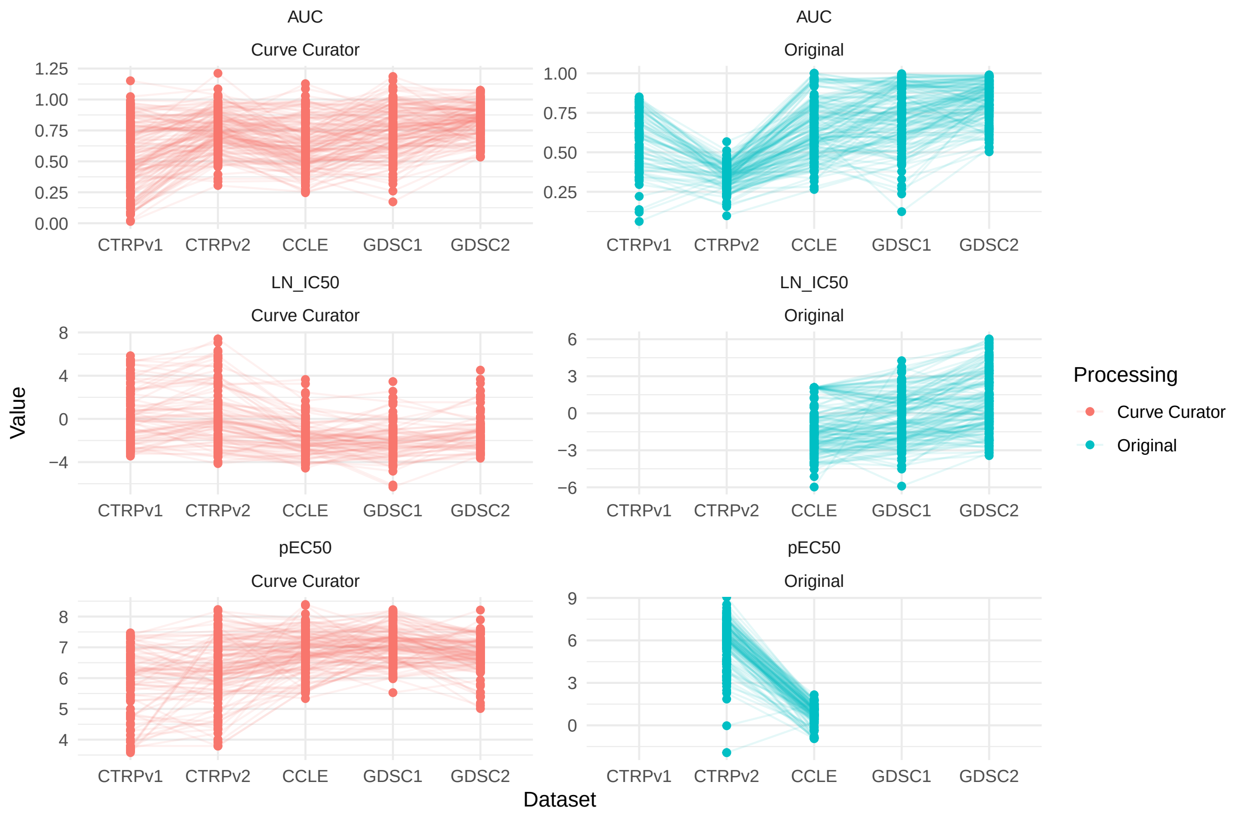


(a) One data point corresponds to one drug-cell line combination. Overall, 148 combinations occurred in all five datasets.

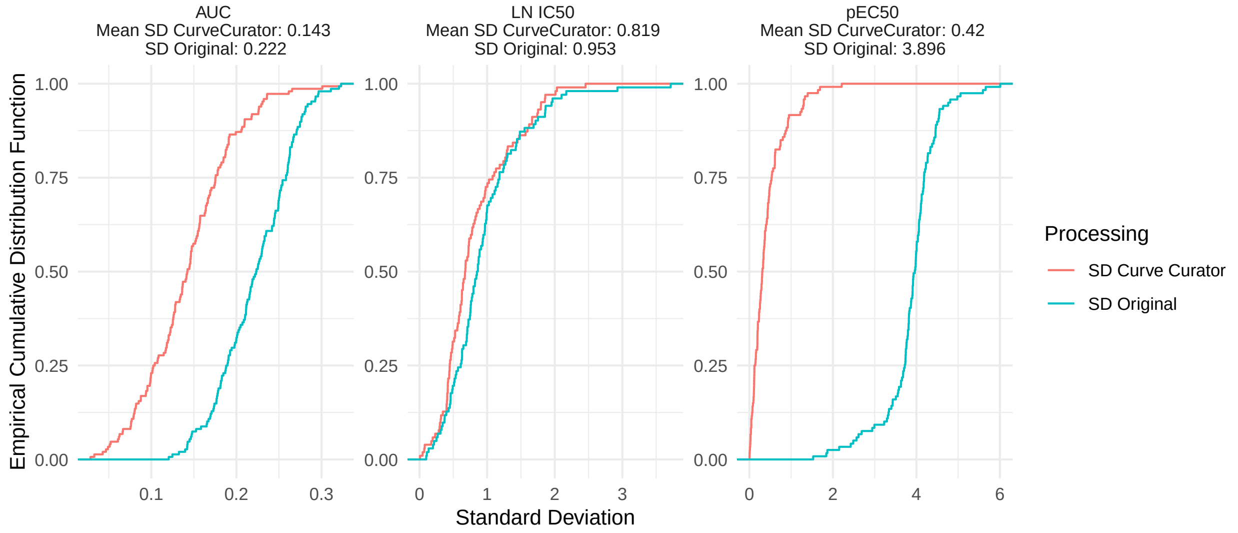


(b) ECDF displaying the standard deviations of the measurements of the same drug-cell line combinations between datasets.

Figure S1: Comparison of response measures in the original screen vs. reprocessed by CurveCurator. Panel b) quantifies how parallel the lines in Panel a) are to the x-axis: If responses roughly agree with each other between datasets, the standard deviation is small. It can be seen that through the uniform preprocessing with CurveCurator, responses agree more with each other.

| | CCLE | CTRPv1 | CTRPv2 | GDSC1 | GDSC2 | |
|---|---|---|---|---|---|---|
| **Nr. of CLs in drug screen** | **503** | **242** | **886** | **970** | **969** | |
| **Modality** | **Overlaps** | | | | | **Nr. of CLs in omics screen** |
| RNAseq gene expr. CCLE | 474 (94%) | 233 (96%) | 820 (93%) | 628 (65%) | 629 (65%) | 1019 |
| Microarray gene expr. GDSC | 382 (76%) | 182 (75%) | 590 (66%) | 943 (97%) | 941 (97%) | 1010 |
| RRBS methylation CCLE | 415 (82%) | 210 (87%) | 733 (83%) | 554 (57%) | 555 (57%) | 842 |
| BeadChip methylation GDSC | 389 (77%) | 184 (76%) | 596 (67%) | 948 (98%) | 947 (98%) | 1025 |
| CNV Cell Model Passports | 369 (73%) | 180 (74%) | 575 (64%) | 951 (98%) | 951 (98%) | 978 |
| Mutation Cell Model Passports | 457 (91%) | 216 (89%) | 759 (85%) | 965 (99%) | 964 (99%) | 1269 |
| DIA Proteomics | 368 (73%) | 177 (73%) | 566 (64%) | 939 (97%) | 944 (97%) | 949 |

Table S2: Coverage of omic screens for the different drug screens employed in this study. The RNAseq data is reprocessed from Ghandi et al. [2019] using the nf-core/rnaseq pipeline. The microarray data and the BeadChip methylation data was downloaded from the GDSC data portal. The RRBS methylation data was downloaded from DepMap. The DIA proteomics screen is from Gonçalves et al. [2022].

| Model / Hyperparameter | Value |
|---|---|
| NaivePredictor | None |
| NaiveDrugMeanPredictor | None |
| NaiveCellLineMeanPredictor | None |
| NaiveTissueMeanPredictor | None |
| NaiveMeanEffectsPredictor | None |
| ElasticNet | l1_ratio [0, 0.5, 1], alpha [1, 0.8, 0.6, 0.4, 0.2, 0.1, 5, 10, 100] |
| SingleDrugElasticNet | l1_ratio [0.2, 0.5, 0.9], alpha [1, 0.8, 0.6, 0.4, 0.2, 0.1, 5, 10, 100] |
| SingleDrugProteomicsElasticNet | l1_ratio [0.2, 0.5, 0.9], alpha [1, 0.8, 0.6, 0.4, 0.2, 0.1, 5, 10, 100] |
| RandomForest | n_estimators [100], max_depth [5, 10, 30], max_samples [0.2], n_jobs [-1], criterion [squared_error] |
| MultiOmicsRandomForest | n_estimators [100], max_depth [5, 10, 30], max_samples [0.2], n_jobs [-1], criterion [squared_error], n_components [100] |
| SVR | kernel [rbf], C [0.001, 0.01, 0.1, 1, 10, 100], epsilon [0.001, 0.01, 0.1, 0.5, 1], max_iter [500] |
| SingleDrugRandomForest | n_estimators [100], max_depth [5, 10, 30], max_samples [0.2], n_jobs [-1], criterion [squared_error] |
| GradientBoosting | max_iter [100], learning_rate [0.1, 0.01], max_depth [5, 10, 30] |
| SimpleNeuralNetwork | dropout_prob: [0.3], units_per_layer: [[32, 16, 8, 4], [128, 64, 32], [64, 64, 32], [1024, 128, 64, 16]], max_epochs: [100] |
| MultiOmicsNeuralNetwork | dropout_prob: [0.3], units_per_layer: [[16, 8, 4], [32, 16, 8, 4], [128, 64, 32], [64, 64, 32]], methylation_pca_components: [100], max_epochs: [100] |

Continued on next page

Table S3: Hyperparameters defined per model. Values are either scalar values or lists of tested values.. If a publication reported only a single set of hyperparameters without documenting a tuning strategy, we fix the configuration to the given values. This prevents possible further implicit data dredging, which can occur when hyperparameters are informally optimized based on performance on the test set instead of on an isolated validation set.

| Model / Hyperparameter | Value |
|---|---|
| MultiOmicsNeuralNetwork | None |
| SRMF | K: 45, lambda_l: 0.01, lambda_d: 0, lambda_c: 0.01, max_iter: 50, seed: 1, n_features: 1036 |
| DIPK | batch_size: [64], lr: [0.0001], heads: [2], fc_layer_num: [3], fc_layer_dim: [[256, 128, 64, 32, 16, 1]], dropout_rate: [0.3], epochs: [100], epochs_autoencoder: [100], patience: [10] |
| MOLIR | mini_batch: [32], h_dim1: [64, 16], h_dim2: [64, 16], h_dim3: [64, 16], learning_rate: [0.01], dropout_rate: [0.5], weight_decay: [0.0001], gamma: [0.5], epochs: [30], margin: [1.5] |
| SuperFELTR | mini_batch: 55, dropout_rate: [0.5], weight_decay: 0.01, out_dim_expr_encoder: 256, out_dim_mutation_encoder: 32, out_dim_cnv_encoder: 64, epochs: 30, margin: 1.0, learning_rate: 0.01 |

### 2 Supplementary Results: Benchmark

Table S4: $lnIC50$ prediction results evaluated per cross-validation (CV) fold, summarized as mean ± standard error. The Pearson per drug/cell line is calculated per model and test mode (CV folds combined). The underlined NaiveMeanEffectsPredictor represents the best possible performance without utilizing any features, by exploiting cell line and drug biases. The normalized metrics are derived by subtracting the predictions of the NaiveMeanEffectsPredictor from the true and predicted values and then recalculating the metric. This predictor is equivalent to the NaiveDrugMeanPredictor and the NaiveCellLineMeanPredictor for LCO/LTO and LDO, respectively. SRMF and SuperFELTR are omitted from LDO as they are not capable of predicting unseen drugs.

| Model | MSE | $R^2$ | $R^2$: normalized | Pearson | Pearson: normalized | Pearson per drug | Pearson per cell line |
|---|---|---|---|---|---|---|---|
| | | | LPO | | | | |
| DIPK | **1.15±0.01** | **0.82±0.00** | **0.32±0.01** | **0.91±0.00** | **0.58±0.01** | **0.53±0.00** | **0.89±0.00** |
| SimpleNeuralNetwork | 1.39±0.02 | 0.78±0.00 | 0.18±0.01 | 0.89±0.00 | 0.44±0.01 | 0.45±0.00 | 0.87±0.00 |
| RandomForest | 1.59±0.01 | 0.75±0.00 | 0.06±0.00 | 0.87±0.00 | 0.29±0.00 | 0.39±0.00 | 0.86±0.00 |
| MultiOmicsRandomForest | 1.63±0.02 | 0.75±0.00 | 0.05±0.00 | 0.87±0.00 | 0.27±0.00 | 0.38±0.00 | 0.86±0.00 |
| MultiOmicsNeuralNetwork | 1.65±0.04 | 0.74±0.01 | 0.03±0.02 | 0.86±0.00 | 0.29±0.02 | 0.39±0.00 | 0.85±0.00 |
| NaiveMeanEffectsPredictor | 1.70±0.01 | 0.74±0.00 | 0.00±0.00 | 0.86±0.00 | 0.00±0.00 | 0.40±0.00 | 0.84±0.00 |
| GradientBoosting | 1.75±0.01 | 0.73±0.00 | -0.03±0.00 | 0.86±0.00 | 0.15±0.00 | 0.42±0.00 | 0.84±0.00 |
| NaiveDrugMeanPredictor | 2.09±0.01 | 0.67±0.00 | -0.23±0.00 | 0.82±0.00 | 0.02±0.00 | -0.21±0.00 | 0.84±0.00 |
| SRMF | 2.77±0.12 | 0.57±0.02 | -0.64±0.07 | 0.81±0.01 | 0.44±0.01 | 0.51±0.00 | 0.80±0.00 |
| ElasticNet | 4.07±0.01 | 0.37±0.00 | -1.41±0.01 | 0.61±0.00 | 0.00±0.00 | 0.43±0.00 | 0.57±0.00 |
| NaiveCellLineMeanPredictor | 6.04±0.02 | 0.06±0.00 | -2.58±0.02 | 0.24±0.00 | 0.00±0.00 | 0.42±0.00 | -0.19±0.00 |
| NaiveTissueMeanPredictor | 6.26±0.02 | 0.03±0.00 | -2.71±0.02 | 0.16±0.00 | 0.02±0.00 | 0.29±0.00 | -0.02±0.00 |
| NaivePredictor | 6.42±0.02 | 0.00±0.00 | -2.81±0.02 | 0.00±0.00 | 0.01±0.00 | 0.00±0.00 | -0.01±0.00 |
| SuperFELTR | 7.10±0.02 | -0.10±0.00 | -3.26±0.03 | 0.28±0.00 | 0.01±0.00 | 0.27±0.00 | 0.28±0.00 |
| | | | LCO | | | | |
| RandomForest | **1.70±0.03** | **0.74±0.01** | **0.19±0.02** | **0.86±0.00** | **0.43±0.02** | **0.32±0.00** | **0.86±0.00** |
| MultiOmicsRandomForest | 1.74±0.02 | 0.73±0.00 | 0.16±0.02 | **0.86±0.00** | 0.41±0.02 | 0.32±0.00 | **0.86±0.00** |
| DIPK | 1.86±0.03 | 0.71±0.01 | 0.11±0.02 | 0.85±0.00 | 0.38±0.02 | 0.30±0.00 | **0.86±0.00** |
| SimpleNeuralNetwork | 1.94±0.03 | 0.70±0.01 | 0.07±0.02 | 0.84±0.00 | 0.35±0.02 | 0.26±0.00 | 0.85±0.00 |
| GradientBoosting | 1.98±0.04 | 0.69±0.01 | 0.05±0.01 | 0.84±0.00 | 0.29±0.01 | 0.27±0.00 | 0.84±0.00 |
| MultiOmicsNeuralNetwork | 2.08±0.04 | 0.68±0.01 | 0.00±0.03 | 0.83±0.00 | 0.28±0.02 | 0.26±0.00 | 0.83±0.00 |
| NaiveMeanEffectsPredictor | 2.09±0.04 | 0.68±0.00 | 0.00±0.00 | 0.82±0.00 | 0.00±0.00 | -0.21±0.00 | 0.84±0.00 |
| SRMF | 2.82±0.26 | 0.56±0.04 | -0.34±0.11 | 0.79±0.00 | 0.09±0.01 | 0.08±0.00 | 0.82±0.00 |
| ElasticNet | 5.89±0.12 | 0.09±0.02 | -1.84±0.08 | 0.44±0.01 | 0.07±0.01 | 0.10±0.00 | 0.54±0.00 |
| NaiveTissueMeanPredictor | 6.28±0.03 | 0.02±0.00 | -2.03±0.05 | 0.15±0.01 | 0.06±0.01 | 0.27±0.00 | — |
| NaivePredictor | 6.42±0.02 | 0.00±0.00 | -2.10±0.05 | 0.00±0.00 | 0.00±0.00 | -0.02±0.00 | — |
| SuperFELTR | 7.15±0.05 | -0.11±0.01 | -2.50±0.08 | 0.32±0.01 | 0.02±0.01 | 0.24±0.00 | 0.37±0.00 |

Continued on next page

Table S5: CONTINUED: $lnIC50$ prediction results evaluated per cross-validation (CV) fold, summarized as mean ± standard error. The Pearson per drug/cell line is calculated per model and test mode (CV folds combined). The underlined NaiveMeanEffectsPredictor represents the best possible performance without utilizing any features, by exploiting cell line and drug biases. The normalized metrics are derived by subtracting the predictions of the NaiveMeanEffectsPredictor from the true and predicted values and then recalculating the metric. This predictor is equivalent to the NaiveDrugMeanPredictor and the NaiveCellLineMeanPredictor for LCO/LTO and LDO, respectively. SRMF and SuperFELTR are omitted from LDO as they are not capable of predicting unseen drugs.

| Model | MSE | $R^2$ | $R^2$: normalized | Pearson | Pearson: normalized | Pearson per drug | Pearson per cell line |
|---|---|---|---|---|---|---|---|
| | | | LTO | | | | |
| RandomForest | **1.80±0.06** | **0.71±0.01** | **0.07±0.01** | **0.85±0.00** | **0.29±0.02** | **0.27±0.00** | **0.85±0.00** |
| MultiOmicsRandomForest | 1.87±0.09 | 0.70±0.01 | 0.05±0.03 | 0.84±0.01 | 0.26±0.03 | 0.24±0.00 | **0.85±0.00** |
| DIPK | 1.99±0.10 | 0.68±0.01 | -0.02±0.03 | 0.83±0.01 | 0.23±0.02 | 0.19±0.00 | 0.84±0.00 |
| GradientBoosting | 2.05±0.10 | 0.67±0.01 | -0.05±0.02 | 0.83±0.00 | 0.14±0.02 | 0.24±0.00 | 0.83±0.00 |
| SimpleNeuralNetwork | 2.12±0.08 | 0.66±0.01 | -0.09±0.02 | 0.82±0.00 | 0.19±0.02 | 0.20±0.00 | 0.83±0.00 |
| NaiveMeanEffectsPredictor | 2.14±0.16 | 0.66±0.02 | -0.09±0.04 | 0.83±0.00 | 0.00±0.00 | -0.39±0.00 | 0.84±0.00 |
| MultiOmicsNeuralNetwork | 2.22±0.13 | 0.65±0.01 | -0.13±0.03 | 0.81±0.01 | 0.14±0.02 | 0.21±0.00 | 0.82±0.00 |
| SRMF | 5.44±0.78 | 0.14±0.11 | -1.75±0.36 | 0.77±0.01 | 0.02±0.01 | -0.01±0.00 | 0.79±0.00 |
| ElasticNet | 6.16±0.75 | 0.04±0.09 | -2.10±0.26 | 0.41±0.01 | 0.04±0.01 | 0.19±0.00 | 0.48±0.00 |
| NaivePredictor | 6.41±0.23 | -0.02±0.01 | -2.29±0.07 | 0.00±0.00 | 0.01±0.02 | -0.25±0.00 | — |
| | | | LDO | | | | |
| MultiOmicsNeuralNetwork | **5.75±0.38** | **0.09±0.03** | **0.02±0.03** | 0.34±0.03 | 0.26±0.04 | 0.33±0.00 | 0.28±0.00 |
| GradientBoosting | 5.86±0.52 | 0.08±0.03 | 0.01±0.03 | 0.29±0.06 | 0.21±0.05 | 0.36±0.00 | 0.24±0.00 |
| ElasticNet | 6.10±0.52 | 0.04±0.01 | -0.03±0.01 | 0.24±0.02 | 0.12±0.02 | 0.32±0.00 | 0.16±0.00 |
| SimpleNeuralNetwork | 5.87±0.34 | 0.06±0.03 | -0.01±0.04 | 0.32±0.04 | 0.25±0.04 | 0.33±0.00 | 0.27±0.00 |
| DIPK | 5.91±0.47 | 0.05±0.06 | -0.02±0.06 | **0.41±0.04** | **0.35±0.05** | **0.42±0.00** | **0.34±0.00** |
| NaiveMeanEffectsPredictor | 6.03±0.53 | 0.05±0.01 | -0.02±0.01 | 0.26±0.01 | 0.00±0.00 | **0.42±0.00** | -0.18±0.00 |
| MultiOmicsRandomForest | 6.23±0.63 | 0.03±0.05 | -0.04±0.06 | 0.30±0.06 | 0.23±0.06 | 0.37±0.00 | 0.23±0.00 |
| NaiveTissueMeanPredictor | 6.28±0.54 | 0.01±0.01 | -0.06±0.01 | 0.16±0.01 | 0.01±0.01 | 0.29±0.00 | -0.12±0.00 |
| NaivePredictor | 6.44±0.54 | -0.02±0.00 | -0.09±0.01 | 0.00±0.00 | 0.01±0.01 | — | -0.12±0.00 |
| RandomForest | 6.63±0.65 | -0.04±0.06 | -0.12±0.06 | 0.26±0.05 | 0.20±0.05 | 0.36±0.00 | 0.20±0.00 |

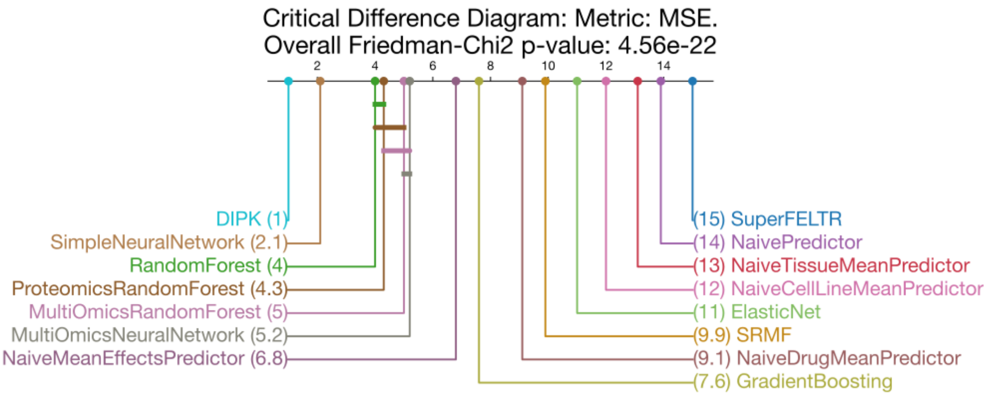


(a) LPO setting.

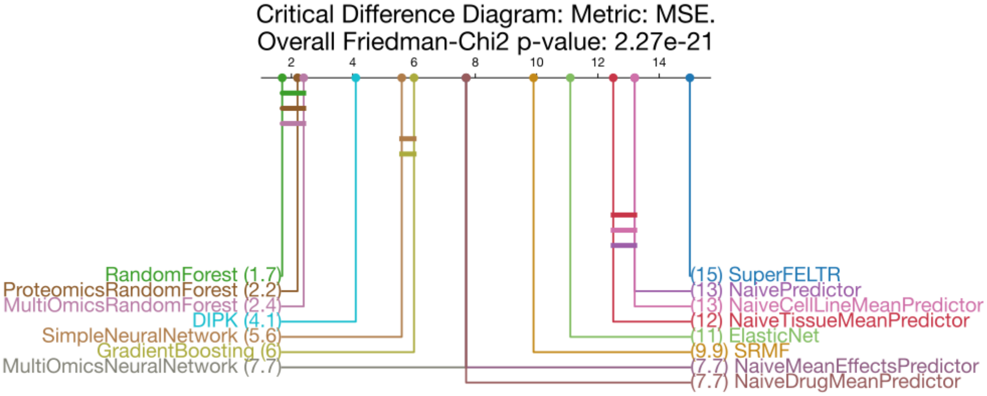


(b) LCO setting.

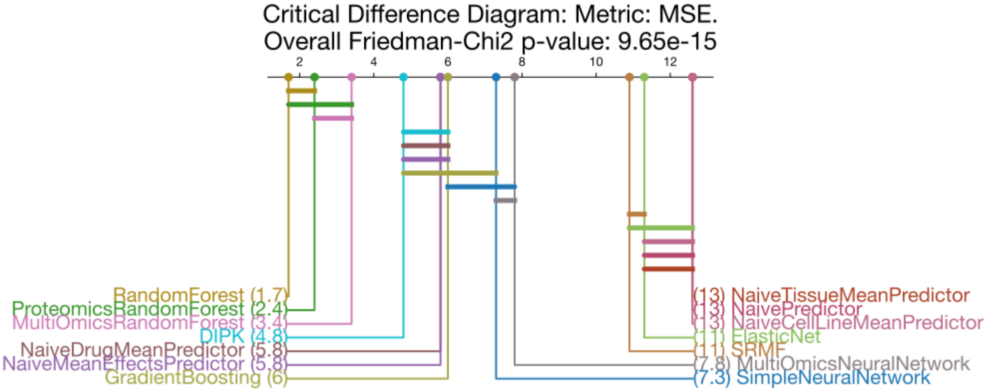


(c) LTO setting.

Figure S2: Critical difference diagram for the LPO (a), LCO (b), and LTO (c) setting using an MSE-based ranking in the cross-validation folds. For each model, we draw an individual horizontal bar, connecting it to all other models from which it does not differ significantly. In the LPO and LCO setting, most differences in model performance are significant.

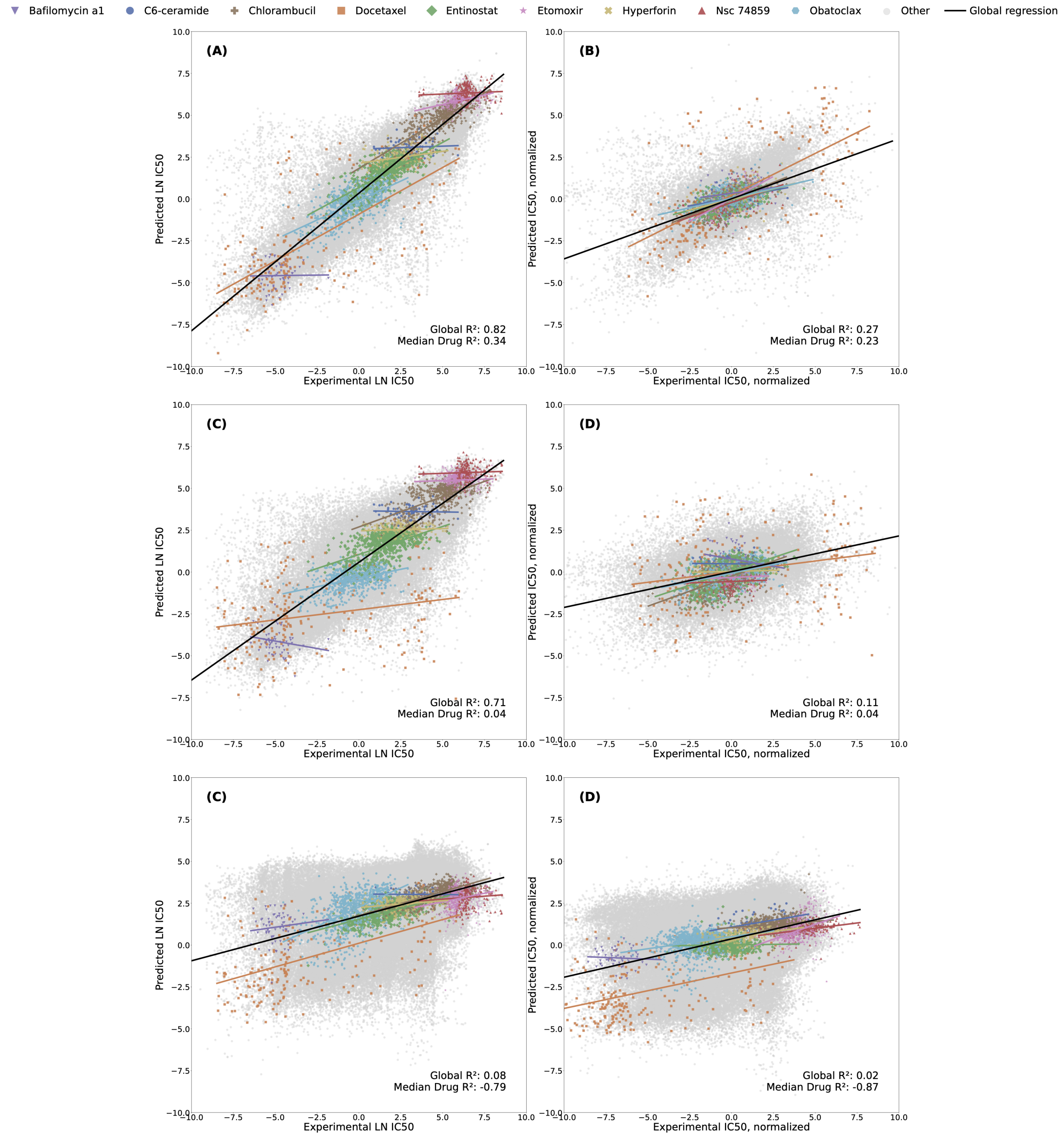


Figure S3: Simpson's paradox in drug response prediction. (A) Predicted log IC50 vs. ground truth for the DIPK model under leave-pair-out cross-validation. The apparent correlation is largely driven by differences in mean drug potency. (B) After subtracting drug and cell line mean effects, only weak signal remains, indicating limited learning of differential response beyond remembering mean cell line and drug responses. (C) Naive coefficients of determinations are lower under leave-cell-line-out cross-validation. (D) However, after normalization with the drug means, the model retains more differential signal compared to LPO. (E), (F) The model can not predict drug responses of unseen drugs.

Table S6: Mean and standard deviation of model runtimes to train and test one LPO split. Note that SuperFELT is a single-drug model so the run time multiplies with the number of drugs for which it is trained.

| Model | Runtime |
|---|---|
| DIPK | 136m 4s ± 22m 20s |
| MultiOmicsNeuralNetwork | 59m 36s ± 32m 25s |
| SimpleNeuralNetwork | 22m 0s ± 7m 18s |
| RandomForest | 11m 33s ± 6m 9s |
| GradientBoosting | 1m 47s ± 0m 28s |
| SRMF | 1m 41s ± 0m 14s |
| ElasticNet | 1m 36s ± 0m 54s |
| NaiveMeanEffectsPredictor | 0m 49s ± 0m 8s |
| SuperFELTR | 0m 41s ± 0m 6s |
| NaiveCellLineMeanPredictor | 0m 36s ± 0m 4s |
| NaiveDrugMeanPredictor | 0m 28s ± 0m 10s |
| NaivePredictor | 0m 10s ± 0m 8s |

#### 2.1 Cross-study prediction on GDSC1, GDSC2

We assessed how model performance is affected when both the input features and the output domain (i.e., drug response measurements) differ from the training data distribution. This scenario applies to GDSC1 and GDSC2, where gene expression and methylation data are primarily available as microarray and Illumina BeadChip measurements, respectively, rather than the RNA-seq and RRBS data used during training. Cross-study performance using RNA-seq and RRBS inputs on these datasets has been previously explored [Xia et al., 2022, Partin et al., 2023]. However, substantial differences exist between the platforms: RMA-normalized microarray expression data is not only limited in feature coverage but also lies on a different scale compared to RNA-seq TPMs (Figure S4), and RRBS CpG clusters do not directly correspond to BeadChip-defined CpG islands. To address this, we mapped RRBS cluster regions to the closest matching BeadChip islands based on maximum overlap, discarding clusters without a suitable match or where a more compatible cluster existed (details in the data preprocessing GitHub repository).

Among all datasets, GDSC1 exhibits the highest cross-study prediction errors, likely due to the additional challenge posed by its use of a different viability assay (Syto60), in contrast to CellTiter-Glo used in all other datasets. This makes generalization to GDSC1 particularly difficult, as it involves both out-of-distribution inputs and a shift in the output distribution. Overall, prediction errors on GDSC1 and GDSC2 are substantially higher than on CTRPv1 and CCLE (as expected), as none of the model weights are tuned to this domain.

Table S7: MSE values  standard errors for all cross-study predictions.

| Model | CTRPv2 | Cross-study: CTRPv1 | Cross-study: CCLE | Cross-study: GDSC1 | Cross-study: GDSC2 |
|---|---|---|---|---|---|
| | | | LPO | | |
| DIPK | **1.15±0.00** | 4.13±0.02 | 5.62±0.04 | 8.99±0.10 | 7.66±0.09 |
| SimpleNeuralNetwork | 1.39±0.01 | 4.05±0.01 | 4.41±0.05 | 11.87±0.19 | 11.64±0.21 |
| RandomForest | 1.59±0.00 | **3.49±0.01** | 3.70±0.01 | 5.88±0.01 | 5.43±0.01 |
| NaiveMeanEffectsPredictor | 1.70±0.00 | 4.72±0.00 | 4.12±0.00 | 6.60±0.00 | 5.79±0.00 |
| GradientBoosting | 1.75±0.00 | 3.73±0.01 | **3.43±0.02** | **5.86±0.03** | **5.35±0.02** |
| NaiveDrugMeanPredictor | 2.09±0.00 | 4.25±0.00 | 3.78±0.00 | 6.45±0.00 | 5.82±0.00 |
| SRMF | 2.77±0.04 | 12.20±0.26 | 15.34±0.18 | 6658.29±3.56 | 5903.23±4.20 |
| ElasticNet | 4.07±0.00 | 6.43±0.04 | 4.43±0.02 | 11.19±0.27 | 11.37±0.27 |
| NaiveCellLineMeanPredictor | 6.04±0.01 | 6.52±0.00 | 7.19±0.01 | 7.84±0.00 | 7.92±0.00 |
| NaivePredictor | 6.42±0.01 | 5.99±0.00 | 6.70±0.01 | 7.70±0.00 | 7.98±0.00 |
| SuperFELTR | 7.10±0.01 | 6.95±0.00 | 6.46±0.01 | 6.92±0.00 | 7.95±0.00 |
| | | | LCO | | |
| DIPK | 1.86±0.01 | 3.26±0.02 | 3.80±0.05 | 7.64±0.08 | 6.41±0.07 |
| SimpleNeuralNetwork | 1.94±0.01 | 3.44±0.01 | 3.62±0.04 | 9.16±0.21 | 8.64±0.19 |
| RandomForest | **1.70±0.01** | **2.93±0.01** | 3.31±0.02 | **5.58±0.02** | **5.10±0.02** |
| NaiveMeanEffectsPredictor | 2.09±0.01 | 3.54±0.01 | 3.37±0.02 | 6.12±0.00 | 5.45±0.00 |
| GradientBoosting | 1.98±0.01 | 3.19±0.01 | **3.22±0.03** | 5.63±0.04 | 5.09±0.04 |
| SRMF | 2.82±0.08 | 7.78±0.28 | 5.16±0.12 | 12530.41±15.87 | 10880.02±15.31 |
| ElasticNet | 5.89±0.04 | 6.29±0.06 | 6.16±0.05 | 11.59±0.62 | 11.77±0.62 |
| NaivePredictor | 6.42±0.01 | 6.18±0.02 | 7.68±0.02 | 7.90±0.00 | 8.46±0.00 |
| SuperFELTR | 7.15±0.02 | 7.01±0.01 | 6.38±0.01 | 6.89±0.00 | 7.92±0.01 |
| | | | LDO | | |
| DIPK | 5.91±0.15 | 6.21±0.05 | 4.80±0.14 | 9.21±0.11 | 8.00±0.13 |
| SimpleNeuralNetwork | **5.87±0.11** | 5.90±0.02 | **3.74±0.15** | 7.29±0.15 | 7.60±0.17 |
| RandomForest | 6.63±0.21 | 6.01±0.08 | 5.59±0.25 | 6.84±0.05 | 6.59±0.09 |
| NaiveMeanEffectsPredictor | 6.03±0.17 | 6.78±0.01 | 5.41±0.23 | 7.40±0.02 | 6.98±0.04 |
| GradientBoosting | 5.86±0.16 | **5.63±0.03** | 4.57±0.20 | **6.45±0.03** | **6.15±0.05** |
| ElasticNet | 6.10±0.16 | 5.90±0.02 | 4.82±0.25 | 6.54±0.02 | 6.34±0.03 |
| NaivePredictor | 6.44±0.17 | 6.36±0.01 | 5.18±0.23 | 7.26±0.02 | 7.04±0.04 |

Table S8: Metrics for cross-study predictions with other response measures (all re-computed with CurveCurator). Note that the MSEs are not comparable because of different scales of lnIC50, pEC50, and AUC.

| Model | CTRPv2 | Cross-study: CTRPv1 | Cross-study: CCLE | Cross-study: GDSC1 | Cross-study: GDSC2 |
|---|---|---|---|---|---|
| | | | LCO lnIC50: MSE | | |
| Random Forest | 1.70±0.11 | 2.94±0.08 | 3.31±0.22 | 5.51±0.16 | 4.99±0.16 |
| Naive Mean Effects Predictor | 2.09±0.11 | 3.54±0.11 | 3.37±0.22 | 6.12±0.02 | 5.45±0.03 |
| | | | LCO AUC: MSE | | |
| RandomForest | 0.012±0.001 | 0.065±0.004 | 0.022±0.001 | 0.047±0.002 | 0.036±0.002 |
| NaiveMeanEffectsPredictor | 0.015±0.001 | 0.069±0.004 | 0.023±0.001 | 0.041±0.000 | 0.024±0.000 |
| | | | LCO pEC50: MSE | | |
| RandomForest | 0.681±0.018 | 0.831±0.02 | 1.283±0.033 | 1.077±0.019 | 1.069±0.017 |
| NaiveMeanEffectsPredictor | 0.701±0.019 | 0.784±0.013 | 1.362±0.033 | 1.293±0.003 | 1.525±0.004 |
| | | | LCO lnIC50: Pearson | | |
| RandomForest | 0.86±0.01 | 0.72±0.01 | 0.74±0.01 | 0.42±0.03 | 0.52±0.02 |
| NaiveMeanEffectsPredictor | 0.82±0.01 | 0.65±0.01 | 0.71±0.02 | 0.44±0.00 | 0.55±0.00 |
| | | | LCO AUC: Pearson | | |
| RandomForest | 0.80±0.01 | 0.48±0.01 | 0.76±0.01 | 0.25±0.02 | 0.29±0.02 |
| NaiveMeanEffectsPredictor | 0.75±0.01 | 0.38±0.01 | 0.73±0.01 | 0.28±0.00 | 0.32±0.00 |
| | | | LCO pEC50: Pearson | | |
| RandomForest | 0.68±0.01 | 0.53±0.01 | 0.24±0.02 | 0.19±0.02 | 0.25±0.01 |
| NaiveMeanEffectsPredictor | 0.67±0.01 | 0.53±0.01 | 0.24±0.01 | 0.29±0.00 | 0.34±0.00 |
| | | | LCO lnIC50: $R^2$ | | |
| RandomForest | 0.74±0.02 | 0.51±0.01 | 0.36±0.04 | 0.12±0.03 | 0.24±0.02 |
| NaiveMeanEffectsPredictor | 0.68±0.02 | 0.41±0.01 | 0.36±0.04 | 0.02±0.00 | 0.17±0.00 |
| | | | LCO AUC: $R^2$ | | |
| RandomForest | 0.64±0.02 | 0.06±0.02 | 0.52±0.02 | -0.07±0.04 | -0.48±0.09 |
| NaiveMeanEffectsPredictor | 0.56±0.02 | 0.00±0.02 | 0.49±0.01 | 0.07±0.00 | -0.01±0.01 |
| | | | LCO pEC50: $R^2$ | | |
| RandomForest | 0.46±0.01 | 0.03±0.03 | -0.57±0.04 | -0.01±0.02 | 0.01±0.02 |
| NaiveMeanEffectsPredictor | 0.45±0.01 | 0.09±0.01 | -0.69±0.05 | -0.21±0.00 | -0.42±0.01 |

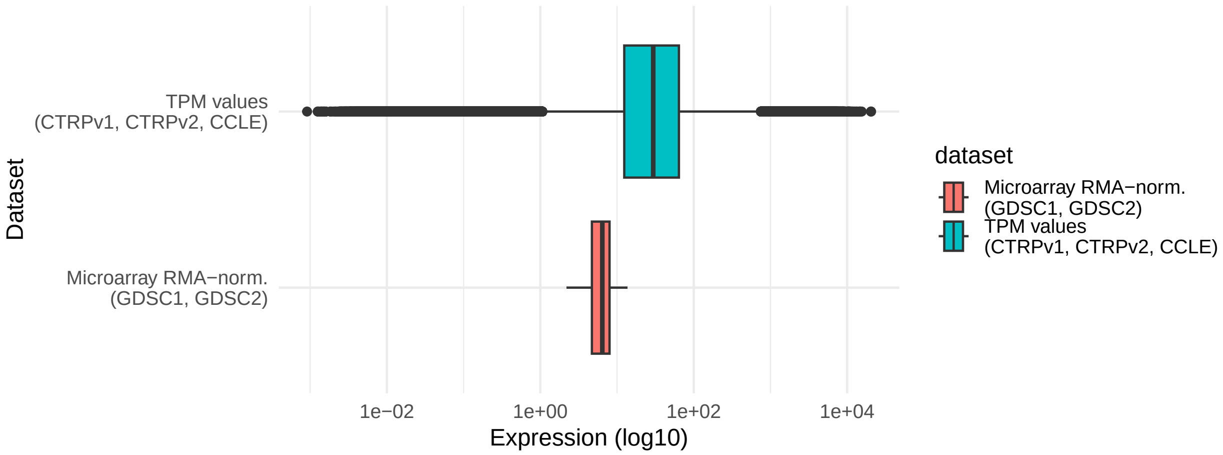


Figure S4: Expression distribution of the microarray vs. the RNA-seq TPM expression values.

#### 2.2 Ablation study

Table S9: Ablation study results of the Multi-OMICS Random Forest and DIPK. One OMIC has been randomized at a time. Perm: a drug/cell line has received a random feature of another drug/cell line. Inv: summary-statistic-invariant randomization.

| Setting | Δ MSE | Δ$R^2$ | Δ Pearson |
|---|---|---|---|
| LPO Multi-OMICS Random Forest | | | |
| CNV (Perm.) | 0.00±0.00 | 0.00±0.00 | 0.00±0.00 |
| CNV (Inv.) | 0.01±0.02 | 0.00±0.00 | 0.00±0.00 |
| **Expression (Perm.)** | 0.15±0.00 | -0.02±0.00 | -0.01±0.00 |
| Expression (Inv.) | 0.13±0.00 | -0.02±0.00 | -0.01±0.00 |
| Methylation (Perm.) | 0.01±0.00 | 0.00±0.00 | 0.00±0.00 |
| Methylation (Inv.) | 0.01±0.00 | 0.00±0.00 | 0.00±0.00 |
| Mutation (Perm.) | 0.00±0.00 | 0.00±0.00 | 0.00±0.00 |
| Mutation (Inv.) | 0.02±0.00 | 0.00±0.00 | 0.00±0.00 |
| Fingerprints (Perm.) | 0.09±0.01 | -0.01±0.00 | -0.01±0.00 |
| Fingerprints (Inv.) | 0.03±0.00 | -0.01±0.00 | 0.00±0.00 |
| LPO DIPK | | | |
| BIONIC (Perm.) | 0.01±0.02 | 0.00±0.00 | 0.00±0.00 |
| BIONIC (Inv.) | 0.00±0.01 | 0.00±0.00 | 0.00±0.00 |
| Expression (Perm.) | 0.06±0.02 | -0.01±0.00 | 0.00±0.00 |
| Expression (Inv.) | -0.04±0.01 | 0.01±0.00 | 0.00±0.00 |
| MolGNet (Perm.) | 0.05±0.02 | -0.01±0.00 | 0.00±0.00 |
| **MolGNet (Inv.)** | 0.06±0.02 | -0.01±0.00 | 0.00±0.00 |
| LCO Multi-OMICS Random Forest | | | |
| CNV (Perm.) | 0.00±0.00 | 0.00±0.00 | 0.00±0.00 |
| CNV (Inv.) | 0.02±0.04 | 0.00±0.00 | 0.00±0.00 |
| **Expression (Perm.)** | 0.20±0.02 | -0.03±0.00 | -0.02±0.00 |
| Expression (Inv.) | 0.17±0.03 | -0.03±0.00 | -0.02±0.00 |
| Methylation (Perm.) | 0.01±0.00 | 0.00±0.00 | 0.00±0.00 |
| Methylation (Inv.) | 0.01±0.04 | 0.00±0.01 | 0.00±0.00 |
| Mutation (Perm.) | 0.00±0.00 | 0.00±0.00 | 0.00±0.00 |
| Mutation (Inv.) | 0.02±0.04 | 0.00±0.00 | 0.00±0.00 |
| Fingerprints (Perm.) | 0.04±0.01 | -0.01±0.00 | 0.00±0.00 |
| Fingerprints (Inv.) | 0.00±0.04 | 0.00±0.00 | 0.00±0.00 |
| LCO DIPK | | | |
| BIONIC (Perm.) | -0.02±0.04 | 0.00±0.01 | 0.00±0.00 |
| BIONIC (Inv.) | 0.02±0.01 | 0.00±0.00 | 0.00±0.00 |
| Expression (Perm.) | 0.15±0.04 | -0.02±0.01 | -0.01±0.00 |
| **Expression (Inv.)** | 0.22±0.03 | -0.03±0.00 | -0.02±0.00 |
| MolGNet (Perm.) | 0.00±0.04 | 0.00±0.01 | 0.00±0.00 |
| MolGNet (Inv.) | 0.05±0.04 | 0.00±0.01 | 0.00±0.00 |
| LDO Multi-OMICS Random Forest | | | |
| CNV (Perm.) | 0.03±0.06 | 0.00±0.01 | 0.00±0.01 |
| CNV (Inv.) | 0.10±0.07 | -0.02±0.01 | -0.01±0.00 |
| Expression (Perm.) | 0.07±0.09 | -0.01±0.02 | -0.01±0.01 |
| Expression (Inv.) | 0.17±0.09 | -0.02±0.01 | -0.03±0.01 |
| Methylation (Perm.) | 0.05±0.07 | -0.01±0.01 | -0.02±0.01 |
| Methylation (Inv.) | 0.14±0.07 | -0.02±0.01 | -0.02±0.01 |
| Mutation (Perm.) | 0.07±0.06 | -0.01±0.01 | -0.01±0.01 |
| Mutation (Inv.) | 0.01±0.09 | -0.01±0.01 | 0.00±0.01 |
| **Fingerprints (Perm.)** | 0.92±0.45 | -0.16±0.07 | -0.15±0.07 |
| Fingerprints (Inv.) | 0.61±0.45 | -0.08±0.06 | -0.11±0.07 |
| LDO DIPK | | | |
| BIONIC (Perm.) | -0.24±0.14 | 0.04±0.02 | 0.00±0.01 |
| BIONIC (Inv.) | -0.09±0.22 | 0.02±0.03 | 0.00±0.02 |
| Expression (Perm.) | -0.29±0.12 | 0.05±0.02 | 0.03±0.01 |
| Expression (Inv.) | -0.08±0.35 | 0.01±0.05 | -0.01±0.03 |
| **MolGNet (Perm.)** | 2.48±0.57 | -0.37±0.08 | -0.30±0.06 |
| MolGNet (Inv.) | 2.21±0.48 | -0.37±0.07 | -0.36±0.06 |

##### 2.3 Proteomics Random Forest

Table S10: Results of the Proteomics Random Forest

| Model | MSE | $R^2$ | $R^2$: normalized | Pearson | Pearson: normalized | Pearson per drug | Pearson per cell line |
|---|---|---|---|---|---|---|---|
| | | | LPO | | | | |
| Random Forest | 1.58±0.04 | 0.75±0.01 | 0.06±0.01 | 0.87±0.00 | 0.30±0.01 | 0.40±0.26 | 0.86±0.08 |
| Proteomics Random Forest | 1.61±0.01 | 0.75±0.00 | 0.05±0.00 | 0.87±0.00 | 0.27±0.00 | 0.38±0.00 | 0.86±0.00 |
| Naive Mean Effects Predictor | 1.70±0.03 | 0.74±0.01 | 0.00±0.00 | 0.86±0.00 | 0.00±0.00 | 0.40±0.25 | 0.84±0.09 |
| | | | LCO | | | | |
| Random Forest | 1.70±0.11 | 0.74±0.02 | 0.19±0.05 | 0.86±0.01 | 0.43±0.05 | 0.33±0.25 | 0.86±0.08 |
| Proteomics Random Forest | 1.73±0.05 | 0.73±0.01 | 0.16±0.01 | 0.86±0.00 | 0.40±0.02 | 0.30±0.00 | 0.86±0.00 |
| Naive Mean Effects Predictor | 2.09±0.11 | 0.68±0.02 | 0.00±0.00 | 0.82±0.01 | 0.00±0.00 | -0.21±0.19 | 0.84±0.08 |
| | | | LTO | | | | |
| Random Forest | 1.82±0.07 | 0.71±0.01 | 0.06±0.02 | 0.85±0.01 | 0.27±0.03 | 0.26±0.00 | 0.85±0.00 |
| Proteomics Random Forest | 1.83±0.07 | 0.71±0.01 | 0.06±0.02 | 0.85±0.01 | 0.26±0.03 | 0.26±0.00 | 0.85±0.00 |
| Naive Mean Effects Predictor | 2.14±0.16 | 0.66±0.02 | -0.09±0.04 | 0.83±0.00 | 0.00±0.00 | -0.39±0.00 | 0.84±0.00 |
| | | | LDO | | | | |
| Naive Mean Effects Predictor | 6.03±1.67 | 0.05±0.02 | -0.02±0.02 | 0.26±0.03 | 0.00±0.00 | 0.42±0.23 | -0.18±0.07 |
| Proteomics Random Forest | 6.14±0.49 | 0.03±0.06 | -0.04±0.06 | 0.32±0.05 | 0.26±0.05 | 0.37±0.00 | 0.25±0.00 |
| Random Forest | 6.58±2.03 | -0.04±0.19 | -0.11±0.20 | 0.27±0.16 | 0.21±0.16 | 0.36±0.26 | 0.21±0.07 |
